# Oligodendrocytes form paranodal bridges that generate chains of myelin sheaths that are vulnerable to degeneration with age

**DOI:** 10.1101/2022.02.16.480718

**Authors:** Cody L. Call, Sarah A. Neely, Jason J. Early, Owen G. James, Lida Zoupi, Anna C. Williams, Siddharthan Chandran, David A. Lyons, Dwight E. Bergles

**Affiliations:** The Solomon Snyder Department of Neuroscience, Johns Hopkins University, Baltimore, Maryland 21205, USA; Centre for Discovery Brain Sciences, University of Edinburgh, Edinburgh EH16 4SB, UK; UK Dementia Research Institute at the University of Edinburgh, Edinburgh EH16 4SB, UK; Centre for Clinical Brain Sciences, University of Edinburgh, Edinburgh EH16 4SB, UK; Euan MacDonald Centre for Motor Neurone Disease Research University of Edinburgh, Edinburgh EH16 4SB, UK; Anne Rowling Regenerative Neurology Clinic, University of Edinburgh, Edinburgh EH16 4SB, UK; Centre for Regenerative Medicine, University of Edinburgh, 5, Little France Drive, Edinburgh, EH16 4UU, UK; Centre for Brain Development and Repair, inStem, Bangalore 560065, India; Johns Hopkins University Kavli Neuroscience Discovery Institute, Baltimore, Maryland 21205, USA

## Abstract

Myelin sheaths in the CNS are generated by the tips of oligodendrocyte processes, which wrap axons to accelerate action potential conduction, provide metabolic support and control excitability. Here we identify a distinct mode of myelination, conserved between zebrafish, mouse and human, in which oligodendrocytes extend myelin along individual axons by linking myelin sheaths across nodes of Ranvier (NoR). By forming thin extensions that cross NoR, which we term paranodal bridges, multiple sheaths can be connected to the soma by a single cytoplasmic process. Extensive *in vivo* live imaging-based analyses, complemented by serial electron microscopic reconstruction of paranodal bridges, revealed that many oligodendrocytes use this strategy to generate longer stretches of myelin along individual axons. In the mouse somatosensory cortex, paranodal bridges were particularly prevalent along the highly branched axons of parvalbumin expressing (PV) interneurons, which enabled oligodendrocytes to extend myelin sheaths around axon bifurcations. Sheaths at the distal ends of these chains of myelin degenerated more frequently in aged mice, suggesting that they may be more vulnerable to the aging brain environment. This previously undescribed and evolutionarily conserved feature of oligodendrocytes extends myelin coverage of individual axons without new oligodendrogenesis, which may reduce metabolic demand and preserve the fidelity of action potential propagation at axon branch points.

## INTRODUCTION

Myelin enables rapid conduction of action potentials within the highly constrained space of the CNS and varies across brain region and neuron type. These distinct patterns of myelin are established as newly generated oligodendrocytes form multilamellar membrane sheaths around a cohort of nearby axons, selected based on diameter, geometry and cellular identity (Almeida et al., 2011; Call and Bergles, 2021; Koudelka et al., 2016; Mayoral et al., 2018; Micheva et al., 2016; Stedehouder et al., 2019, 2018, 2017; Zonouzi et al., 2019). Despite the extensive axonal territory available to oligodendrocytes during development, the length of individual myelin sheaths rarely exceeds 150 µm in the brain, suggesting that sheath extension is tightly constrained by both cell intrinsic and extrinsic mechanisms (Bechler et al., 2015; Chong et al., 2012; Stedehouder et al., 2019). Indeed, oligodendrocytes exposed to long unbranched substrates *in vitro*, such as artificial nanofibers or dorsal root ganglion neurons, form sheaths that are comparable in length to those observed *in vivo* (Bechler et al., 2015). After a period of initial remodeling, individual myelin sheaths are extremely stable, allowing long-term control of action potential conduction (Lang and Rosenbluth, 2003; Micheva et al., 2021; Seidl and Rubel, 2016), metabolic support (Lee et al., 2012; Morrison et al., 2015; Philips et al., 2021; Rinholm et al., 2011; Saab et al., 2013) and axonal excitability (Hamada and Kole, 2015; Larson et al., 2018; Schirmer et al., 2018), suggesting that the precise location of myelin shapes the functional characteristics of neuronal circuits. However, the mechanisms that control the positioning of myelin along individual axons remain to be defined.

The initial phase of myelination is rapid, promiscuous, and error-prone, resulting in the removal of nascent sheaths from unsuitable targets (Czopka et al., 2013; Hughes and Appel, 2020; Orthmann-Murphy et al., 2020), with subsequent longitudinal extension of sheaths on appropriate axons eventually supporting conduction speeds with sub-millisecond precision (Seidl et al., 2014; Seidl and Rubel, 2016). At these sites of interaction along axons, newly generated oligodendrocytes form highly dynamic, nascent sheaths that are remodeled on the timescale of minutes (Czopka et al., 2013; Haber et al., 2009; Ioannidou et al., 2012). Despite initiating wrapping, oligodendrocytes also extend filopodia from the outermost sheath membrane (Haber et al., 2009; Hardy and Friedrich, 1996; Toth et al., 2021), suggesting that their processes continue to search for suitable targets even after initial sheath formation. However, the biological significance of these persistent dynamics and their role in generating the diverse patterns of myelination that exist in the adult CNS have not been established.

Individual axons in the CNS often undergo extensive branching to enable innervation of distinct brain regions and diverse targets within the terminal field. In particular, inhibitory interneurons exhibit prolific axonal branching, allowing the relatively small number of these neurons to synchronize the large groups of excitatory neurons needed to elicit large scale rhythmic activity that occurs during discrete brain states (Cardin et al., 2009; Cobb et al., 1995; Klausberger et al., 2004; Ratnadurai-Giridharann et al., 2015; Wang and Buzsáki, 1996).

Although projecting primarily locally, the highly branched axons of parvalbumin-expressing (PV) interneurons are extensively myelinated, speeding feedback inhibition to prevent runaway excitation and maintain time of arrival within distributed networks. Alterations in PV interneuron myelination have been observed in neurodegenerative diseases and psychiatric conditions, such as schizophrenia (Stedehouder and Kushner, 2017), highlighting the importance of establishing and maintaining myelination patterns on these inhibitory cells. However, the extensive branching of their axons presents challenges for myelination, as axon collateral formation interrupts sheath extension, particularly given that oligodendrocytes typically extend few primary cytoplasmic processes that exhibit limited branching (Czopka et al., 2013; Murtie et al., 2007; Orthmann-Murphy et al., 2020), constraining the formation of multiple internodes on each axon. The mechanisms that oligodendrocytes use to overcome these structural constraints to ensure myelination of highly branched axons in the CNS are not known.

Using a combination of longitudinal *in vivo* imaging, high resolution confocal microscopy, and volumetric serial electron microscopy (EM), we discovered that oligodendrocyte processes frequently form a “paranodal bridge,” an extension of the outer tongue of a myelin sheath, to link sheaths across nodes of Ranvier (NoR). We show that this mode of myelin sheath formation is highly conserved, existing within the developing zebrafish spinal cord, the mouse cerebral cortex, and in human oligodendrocytes within both pluripotent stem cell-derived organoids and postmortem human cortex. Paranodal bridges were formed by many oligodendrocytes examined in these species, indicating that they contribute substantially to overall myelin patterning. In layer II/III of the mouse cortex, paranodal bridges were observed at remarkably high frequency along axons of PV interneurons, which allowed myelin sheaths to extend across axon branch points. Although allowing additional sheath formation without oligodendrogenesis, the distal myelin sheaths connected by a paranodal bridge degenerated more frequently than proximal sheaths in aged mice. Together, these findings reveal a previously undescribed mode of myelin sheath generation that allows continuous myelination of axons with highly branched arbors. The reliance on production of distal sheaths through narrow paranodal bridges may render these myelin segments more vulnerable to degeneration with disease and aging.

## RESULTS

### Paranodal bridges link adjacent myelin sheaths in the cerebral cortex

*In vivo* two photon imaging of the somatosensory cortex in adult *Mobp-EGFP* mice, in which it is possible to visualize the complete morphology of individual oligodendrocytes, including their somata, cytoplasmic processes and full complement of myelin sheaths (Hughes et al., 2018), revealed that some sheaths appeared to be isolated, without an apparent cytoplasmic process linking it to a soma (Figure 1A-C, orange sheaths). The high stability of oligodendrocytes in the cortex (Hill et al., 2018; Hughes et al., 2018; Tripathi et al., 2017; Yeung et al., 2014), and the persistence of these apparently isolated sheaths over many weeks of imaging (Figure 1C) suggested that they were unlikely to be stranded myelin internodes from oligodendrocytes that had degenerated. Inspection of the NoR gaps separating these apparently isolated sheaths from neighboring, cytoplasmic process-bearing sheaths, revealed that in each instance there was a thin bridge of EGFP-containing cytoplasm across the NoR linking the two sheaths (Figure 1D), in contrast to typical neighboring sheaths that exhibited no cytoplasmic continuity (Figure 1E). Although unexpected, these structures were not rare in the cortex. Complete reconstructions of oligodendrocytes revealed that many formed multiple paranodal bridges, with up to 20% of their sheaths connected by these structures (range: 0–11 sheaths; average: 7 ± 1%, n = 34 cells from 15 mice) (Figure 1F, blue sheaths). We hypothesized that these thin cytoplasmic connections serve as bridges to connect the distal sheath to the rest of the oligodendrocyte.

**Figure 1.**
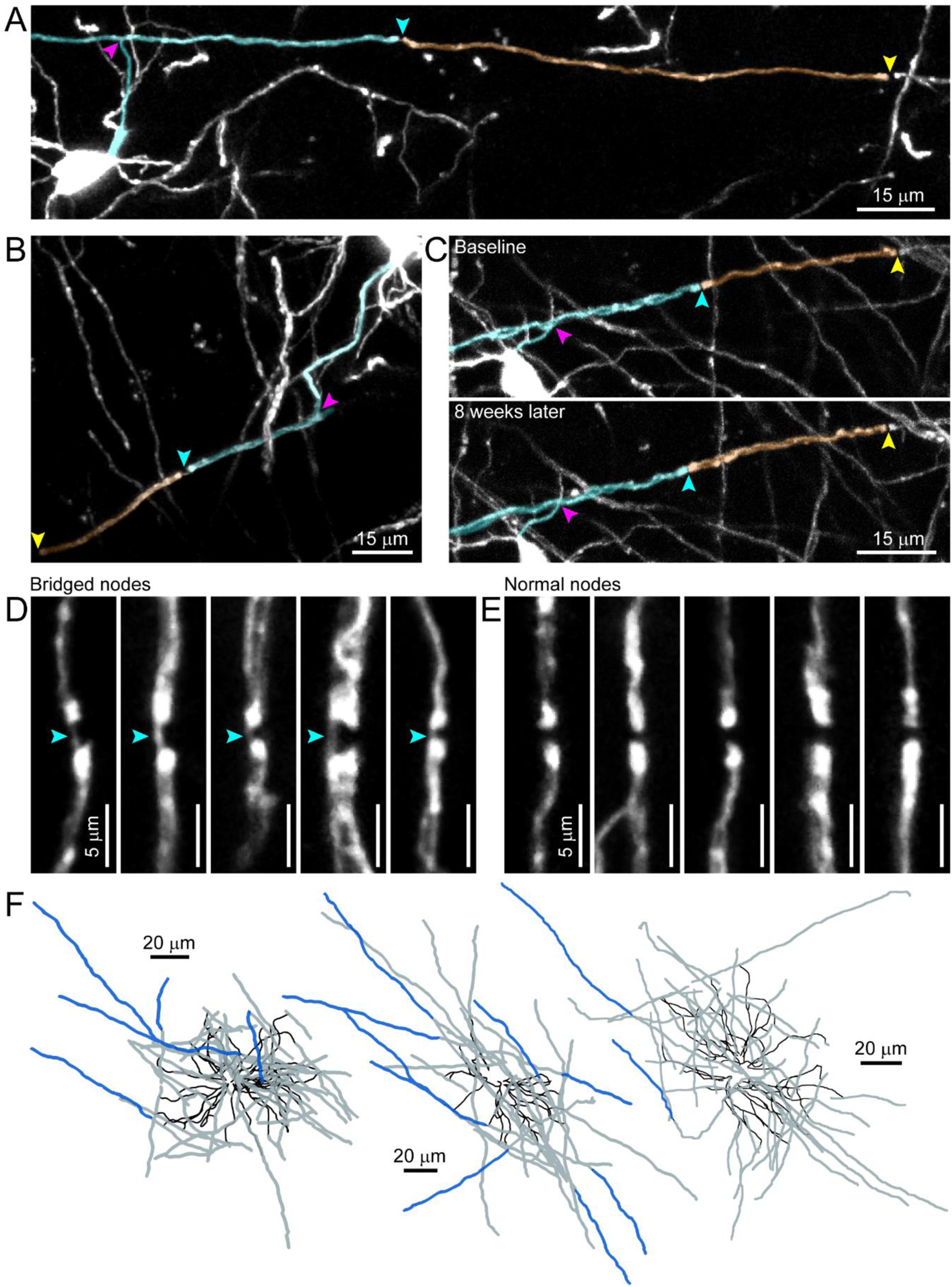
Paranodal bridges in the mouse cerebral cortex. (A) A mouse cortical oligodendrocyte imaged *in vivo* sends a cytoplasmic process to a sheath (magenta arrowhead), which forms a node with an adjacent sheath *via* paranodal bridge (cyan arrowhead). This neighboring sheath terminates at the next node (yellow arrowhead), but has no connecting cytoplasmic process.
(B) Another oligodendrocyte sends a cytoplasmic process to a sheath (magenta arrowhead), which forms a node *via* paranodal bridge (cyan arrowhead) to a secondary sheath (second paranode, yellow arrowhead).
(C) An oligodendrocyte forms a pair of sheaths connected by paranodal bridge (cyan arrowhead) as in *A* and *B* as observed at a baseline imaging timepoint. Eight weeks later (bottom panel), the two sheaths and bridged node remain in nearly identical positions. Magenta arrowhead: cytoplasmic process intersection; yellow arrowhead: distal paranode of bridged sheath.
(D) Five examples of bridged nodes of Ranvier from different cells, with arrowheads indicating paranodal bridge.
(D) Five examples of unbridged nodes of Ranvier from different cells.
(E) Three examples of fully reconstructed cortical oligodendrocyte morphologies imaged *in vivo*. Black: cytoplasmic processes; gray: sheaths connected directly by cytoplasmic process; blue: sheaths connected *via* paranodal bridge. Cell bodies not shown.

To define the structure of paranodal bridges, we immunostained the cerebral cortex of *Mobp-EGFP* mice with myelin basic protein (MBP) and the paranodal protein CASPR and imaged NoR at high resolution. Similar to that observed *in vivo*, thin EGFP+ cytoplasmic extensions were often visible connecting CASPR immunopositive paranodes (Figure 2A,B). In some cases, three sheaths were linked together by two sequential paranodal bridges, in which the center sheath formed paranodal bridges with both neighboring sheaths (Figure 2A,B). In these instances, the “anchoring” sheath (connected to the soma by a cytoplasmic process) could either be the first or second in the chain. Bridged NoR were otherwise indistinguishable from those that lacked paranodal bridges, as they were βIV-spectrin immunopositive (Figure 2C), flanked by CASPR immunoreactivity (Figure 2D), and consistently MBP immunonegative (Figure 2E).

**Figure 2.**
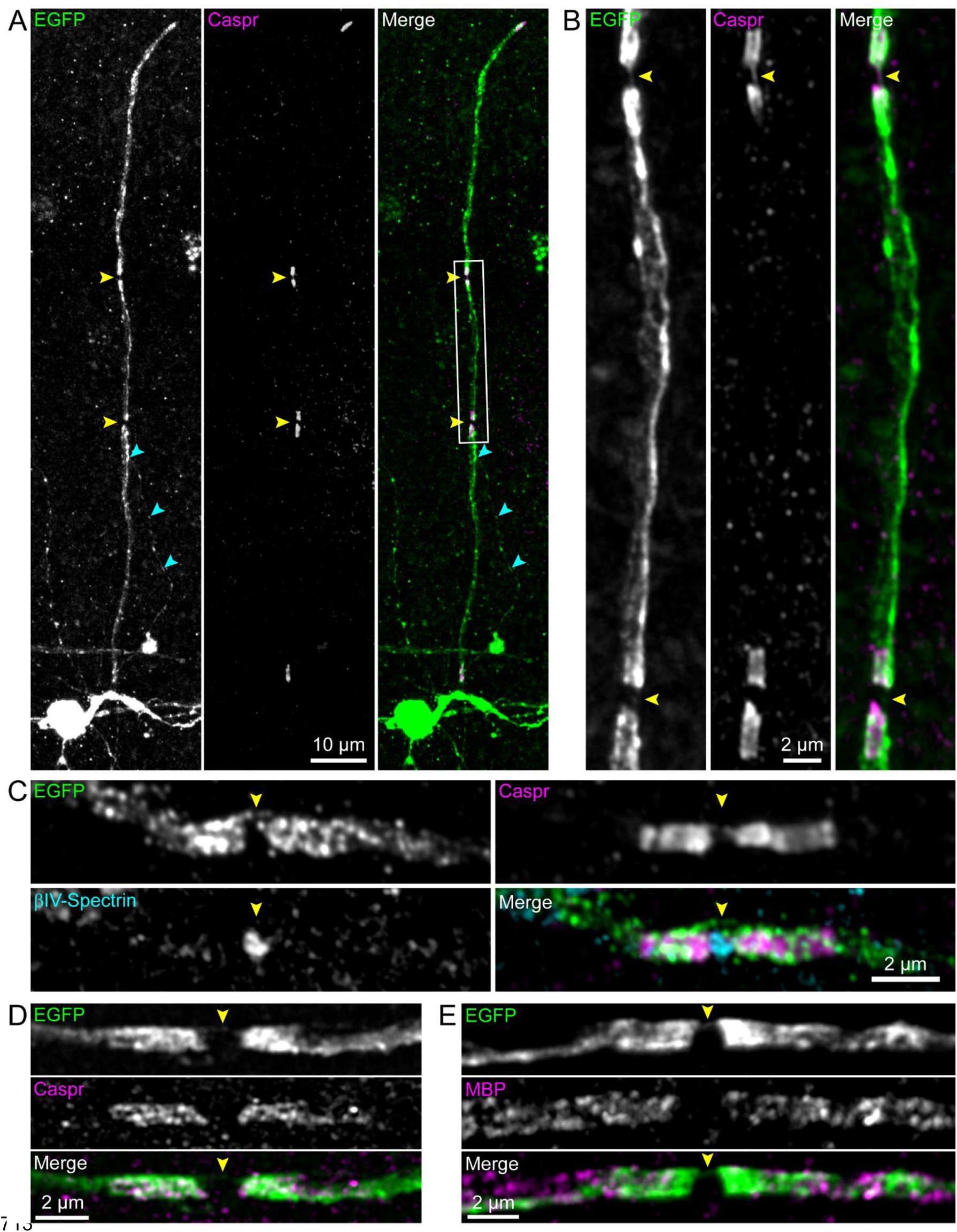
Nodes with paranodal bridges express Caspr, βIV-spectrin, and lack MBP. (A) An individual oligodendrocyte generating a chain of three sheaths linked by two paranodal bridges (yellow arrowheads) imaged from a cortical flatmount of an *Mobp-EGFP* adult mouse, immunostained for EGFP (left/green) and Caspr (center/magenta). Cyan arrowheads indicate cytoplasmic process connecting the cell body to the anchoring sheath.
(B) Magnified view of the white box in *A*.
(C) A magnified view of a node with a paranodal bridge (yellow arrowhead), immunostained for Caspr and βIV-spectrin.
(D) A second example of a node with a paranodal bridge (yellow arrowhead), immunostained for Caspr.
(E) A third example of a node with a paranodal bridge (yellow arrowhead), immunostained for MBP.

To examine the distribution of paranodal bridges among the complement of sheaths formed, we reconstructed the complete morphology of individual oligodendrocytes imaged *in vivo*. This analysis revealed that bridged sheaths typically constituted the most distal internodes, allowing oligodendrocytes to extend myelin further than the reach of typical sheaths; as a result, oligodendrocytes with bridged sheaths had slightly wider territories than those without bridged sheaths (Supplemental Figure 1A-C, Figure 1F). This expansion of myelin territory was particularly apparent in sparsely myelinated regions, such as the temporal association area of the cerebral cortex. In these regions, oligodendrocyte somata are spaced several hundred micrometers apart (Supplemental Figure 2A,B), presenting challenges for establishing continuously myelinated segments. In this environment, chains of sheaths were visible extending between adjacent oligodendrocytes to form NoR (Supplemental Figure 2A,B), with paranodal bridges connecting the center-most sheath (Supplemental Figure 2C). Thus, paranodal bridge formation allows oligodendrocytes to increase the length of axon myelinated without oligodendrogenesis.

### Myelin sheaths connected by paranodal bridges have typical dimensions

To define the features of these distal myelin sheaths relative to other myelin segments, we quantified their properties using *in vivo* two photon imaging of the somatosensory cortex in young adult *Mobp-EGFP* mice. Due to the extended period of oligodendrogenesis in the cortex, it was possible to compare sheath structure from both pre-existing and newly formed oligodendrocytes. Sheaths were defined as non-bridged, anchoring (a sheath containing a typical oligodendrocyte process as well as a cytoplasmic process connected to a bridged sheath), or bridged (a sheath with only a cytoplasmic process coming from another sheath). We refer to a combined unit of an anchoring sheath plus bridged sheath(s) as a bridge chain (Figure 3A). Oligodendrocytes exhibited remarkable diversity in myelin organization in the somatosensory cortex, with bridged sheath content ranging from 0–11 (average: 3 ± 0.5 bridged sheaths per cell). Only 2/34 oligodendrocytes in this region analyzed did not establish bridged sheaths, further highlighting the high incidence of this phenomenon in the cerebral cortex.

**Figure 3.**
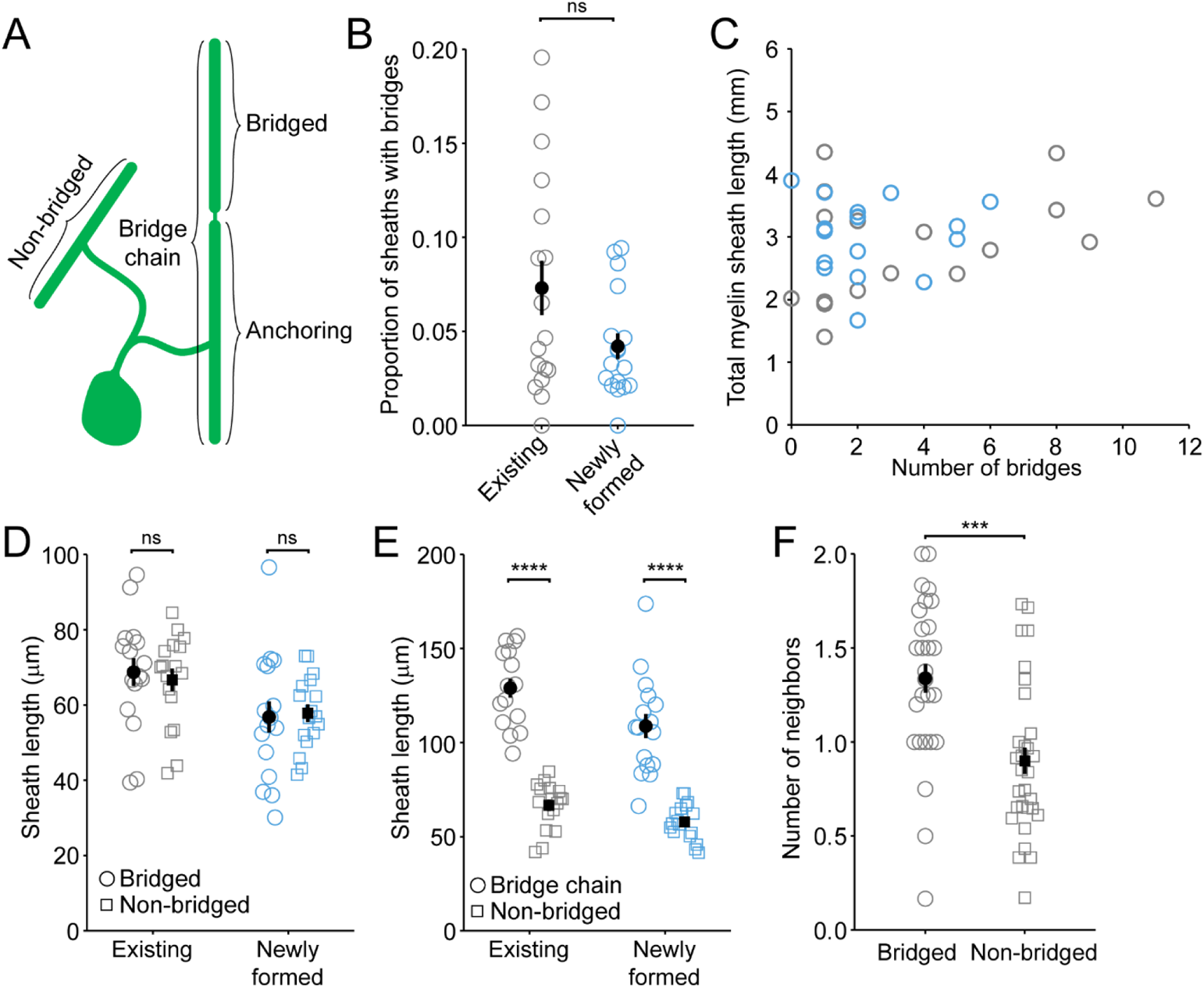
Paranodal bridges are a common feature of oligodendrocytes and link sheaths of typical lengths. (A) Schematic depicting the categories of sheaths examined in this Figure. “Non-bridged” sheaths are canonical sheaths connected directly *via* cytoplasmic process to the cell body. “Bridged” sheaths are connected only *via* paranodal bridge from an “anchoring” sheath connected directly by a cytoplasmic process to the cell body. “Bridge chains” consist of the anchoring sheath and the sheath(s) connected to it *via* paranodal bridge(s).
(B) Bridged sheaths make up a similar proportion of the total cohort of sheaths for individual oligodendrocytes generated prior to 8 weeks of age (existing) and those generated later during *in vivo* imaging time courses (Kruskal-Wallis ANOVA, p = 0.20).
(C) The production of bridged sheaths is not correlated with total cell size (combined length of all sheaths). existing: R^2^ = 0.16, p = 0.11; control: R^2^ = 2.8 × 10^−5^, p = 0.98.
(D) Bridged sheaths (circles) have lengths similar to non-bridged sheaths (squares) across in existing cells, and in newly generated cells (blue) (p = 0.46; Kruskal-Wallis one-way ANOVA).
(E) The combined length of all sheaths connected in a bridge chain is significantly greater than the length of non-bridged sheaths (existing: p = 2.8 × 10^−11^; newly formed: p = 4.1 × 10^−8^, paired two-sample t-tests with Bonferroni correction for multiple comparisons).
(F) Bridged sheaths have more neighbors than non-bridged sheaths (Data combined for existing and newly formed oligodendrocytes. Existing: p = 2.7 × 10^−4^, two-tailed t-test). ***p *<* 0.001; ****p *<* 0.0001; ns, not significant.

Although there was a trend for pre-existing cells to have more bridged sheaths than those generated during the imaging period, this difference was not statistically significant (p = 0.20 Kruskal-Wallis, Figure 3B), indicating that this phenomenon is not limited to early developmental periods. The presence of paranodal bridges did not increase the total myelin sheath length of an oligodendrocyte (number of bridges: 3 ± 0.5; total length: 2.96 ± 0.13 mm; R = 0.26, p = 0.14) (Figure 3C), consistent with previous studies indicating that oligodendrocytes tightly limit their total myelin output (Almeida et al., 2018, 2011; Bacmeister et al., 2020; Chong et al., 2012; Orthmann-Murphy et al., 2020). Sheaths connected via paranodal bridges were also similar in length to all other sheaths (Figure 3D); thus, the combined length of the “chain” of sheaths connected by paranodal bridge(s) was approximately twice the length of a non-bridged sheath (Figure 3E).

Axons in the cerebral cortex vary markedly in their propensity for myelination, reflecting the influence of cell identity, axon diameter and signaling interactions. To determine if extension of myelin through paranodal bridges is more likely to occur on highly *versus* sparsely myelinated axons, we examined whether bridge chains formed NoR with other myelin sheaths or existed as an isolated patch without nearby myelination. Comparing the average number of neighboring sheaths revealed that bridged sheaths were more likely to form NoR with a non-bridged sheath on the other side (i.e. have >1 neighboring sheaths) (Figure 3F) (number of neighbors: bridged, 1.3 ± 0.08; non-bridged, 0.90 ± 0.07; p = 2.7 × 10^−4^, two-tailed t-test). Thus, the properties of axons that increase the likelihood of myelination also promote the formation of chains of bridged myelin sheaths.

### Paranodal bridges are evolutionarily conserved

Oligodendrocyte maturation and myelin formation in the developing zebrafish spinal cord closely resembles that observed in the mouse cerebral cortex (Czopka et al., 2013; Hughes et al., 2013; Kirby et al., 2006; Marisca et al., 2020; Orthmann-Murphy et al., 2020; Snaidero et al., 2014). To determine if paranodal bridges are used to establish myelin in other species, we examined sheaths formed by individual oligodendrocytes in the larval zebrafish spinal cord (4 days post fertilization, dpf), by expressing membrane anchored EGFP (EGFP-CAAX) under control of the myelin basic protein (*mbp*) promoter. *In vivo* analysis of these sparsely labeled oligodendrocytes revealed that 31% of oligodendrocytes (n = 13/42 oligodendrocytes, 41 zebrafish) exhibited paranodal bridges, visible as a narrowing of the EGFP-labeled myelin membrane between adjacent myelin sheaths (Figure 4A,B). Bridged sheaths comprised 15 ± 3% of all sheaths in this region, and were slightly shorter on average than non-bridged sheaths at 4 dpf (non-bridged sheaths: 29.6 ± 2.3 µm; bridged sheaths: 21.2 ± 2.4 µm); nonetheless, the total length of linked sheaths that formed a myelin unit was approximately twice that of a normal sheath (total myelin bridge: 52.2 ± 4.4 µm; repeated measures one-way ANOVA with Tukey’s correction, n = 13 oligodendrocytes) (Figure 4C), comparable to that observed in mouse.

**Figure 4.**
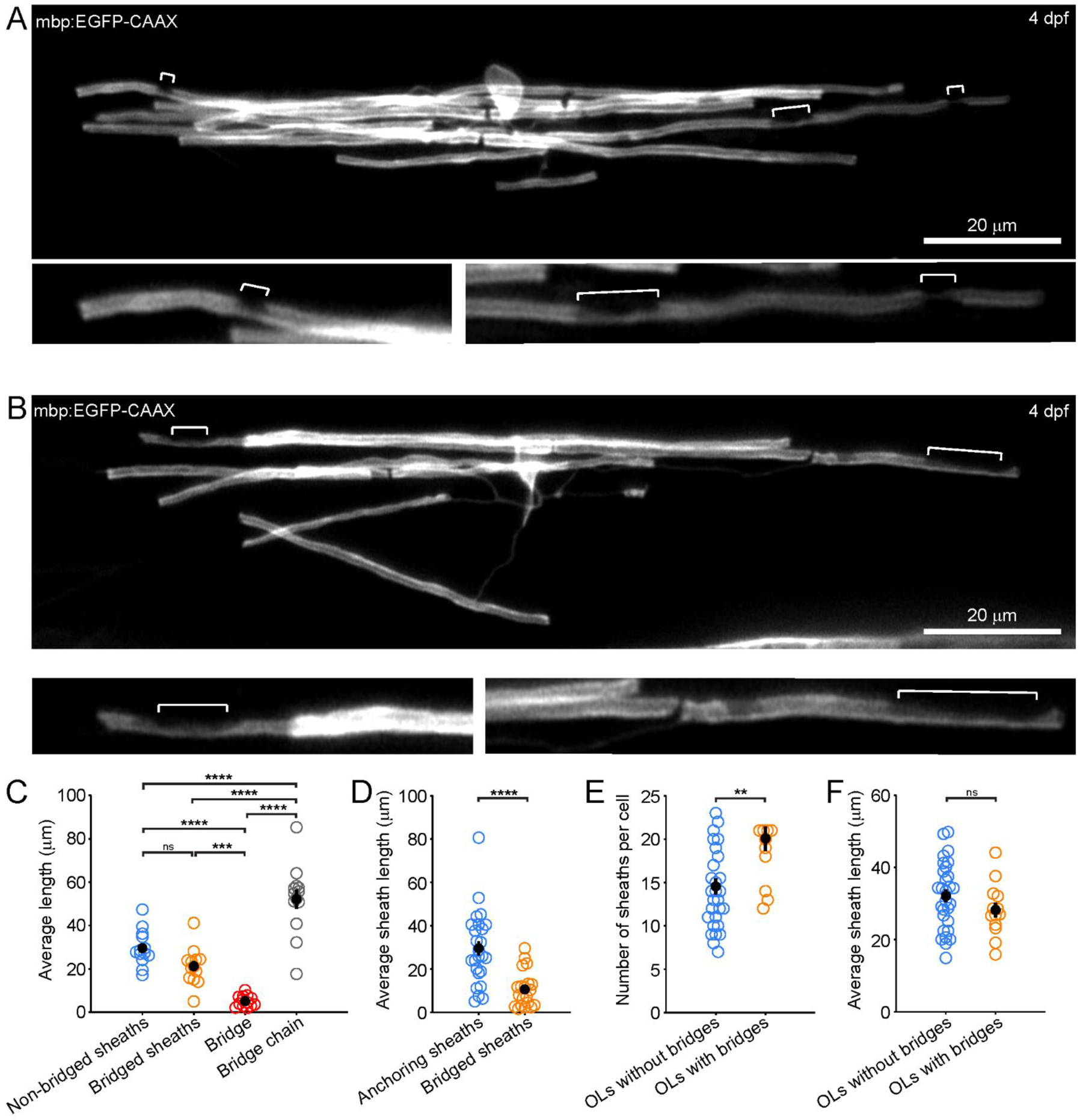
Mature oligodendrocytes have paranodal bridges bridges in the zebrafish spinal cord at 4 dpf. (A) Top panel: confocal image of a single oligodendrocyte at 4 days post fertilisation with paranodal bridges, highlighted with white brackets. Bottom panels: enlarged examples of highlighted bridged nodes.
(B) Top panel: confocal image of a single oligodendrocyte at 4 days post fertilisation with paranodal bridges, highlighted with white brackets. Bottom panels: enlarged examples of highlighted bridged nodes.
(C) Average lengths of non-bridged sheaths (blue, 25.9 ± 1.5 µm), bridged sheaths (orange, 21.2 ± 2.8 µm), paranodal bridges (red, 5.3 ± 0.8 µm), and bridged chains (gray, 49.5 ± 5.7 µm). Repeated measures one-way ANOVA with Tukey’s correction. n = 14 oligodendrocytes.
(D) Quantification of the sheath length of anchoring sheaths (blue, 29.6 ± 3.4 µm) versus bridged sheaths (orange, 10.6 ± 1.9 µm) p = 3.9 × 10^−5^, unpaired two-tailed t-test. n = 25 anchoring sheaths (where 3 bridged sheaths were joined the 2 innermost sheaths were counted as anchoring sheaths) and n = 14 bridged sheaths (from 13 oligodendrocytes with bridged sheaths).
(E) Quantification of the number of myelin sheaths produced per oligodendrocyte in oligodendrocytes without bridges (orange, mean = 15 ± 1 sheaths) versus oligodendrocytes with bridges (blue, 20 ± 1 sheaths) p = 0.0023, unpaired two-tailed t-test. n = 29 oligodendrocytes without bridges and n = 13 oligodendrocytes with bridges.
(F) Quantification of the average myelin sheath length produced per oligodendrocyte in oligodendrocytes without bridges (blue, 32.1 ± 1.7 µm) versus oligodendrocytes with bridges (orange, 28.2 ± 2.1 µm) p = 0.19, unpaired two-tailed t-test. n = 29 oligodendrocytes without bridges and n = 13 oligodendrocytes with bridges. **p *<* 0.01; ***p *<* 0.001; ****p *<* 0.0001; ns, not significant.

However, in zebrafish, anchoring sheaths were typically longer than the bridged sheath(s) (anchoring sheaths: 29.6 ± 3.4 µm; bridged sheaths: 10.6 ± 1.9 µm; p = 3.9 × 10^−5^, unpaired two-tailed t-test) (Figure 4D), and oligodendrocytes that formed paranodal bridges had more sheaths/cell (Figure 4E), but similar average sheath lengths (Figure 4F). These differences may simply reflect distinct features of target axons in the mouse cortex and the zebrafish spinal cord, differences in the relative maturity of oligodendrocytes analysed, or subtle divergence between species.

To determine if human oligodendrocytes also link myelin sheaths through paranodal bridges, we examined the structure of oligodendrocytes in human induced pluripotent stem cell (iPSC)-derived myelinating organoids (“myelinoids”), generated using a previously described spinal cord patterning strategy (James et al., 2021). These myelinoids exhibit formation of MBP expressing oligodendrocytes within 59 days of culture, followed by robust myelination of axons by day 133, with fully compacted sheaths and morphologies consistent with human oligodendrocytes *in vivo* (James et al., 2021). Full reconstructions of oligodendrocytes in myelinoids revealed that many generated paranodal bridges (34%, 53/155 across all myelinoids) (Figure 5A-G). Similar to zebrafish, oligodendrocytes that formed paranodal bridges in myelinoids were more likely to have greater numbers of sheaths than those without bridges (without bridges: 8 ± 1 sheaths; with bridges: 11 ± 1 sheaths; p = 0.0027, generalized linear mixed model) (Figure 5H), with overall average sheath lengths similar for oligodendrocytes with bridged and without bridged sheaths (without bridges: 110 ± 5 µm; with bridges: 103 ± 6 µm; p > 0.05, linear mixed model) (Figure 5I). Also, similar to observations in zebrafish, bridged sheaths in human myelinoids tended to be shorter on average (76 ± 7 µm) compared to both non-bridged (108 ± 4 µm) and anchoring sheaths (112 ± 9 µm, p < 0.001, generalized linear mixed model) (Figure 5J). Moreover, the cumulative length of a bridged chain was significantly longer than individual sheaths (194 ± 16 µm, p < 0.001, generalized linear mixed model) (Figure 5J). In addition, we observed that oligodendrocytes in human organoids frequently extended paranodal bridge-like processes that formed myelin sheaths on neighboring axons, a phenomenon also observed in the mouse cortex (myelinoids: 2/51 bridges; mice: 5/127 bridges) (Supplemental Figure 3A,B), suggesting that process outgrowth can result in bridged sheath formation with other nearby axons. Variations in the structure of bridged sheaths may reflect differences in the properties of the recipient axons, the myelinating environment, or the properties of the oligodendrocytes, which arise from distinct progenitor pools (Cai et al., 2005; Fogarty et al., 2005; Kessaris et al., 2006; Vallstedt et al., 2005).

**Figure 5.**
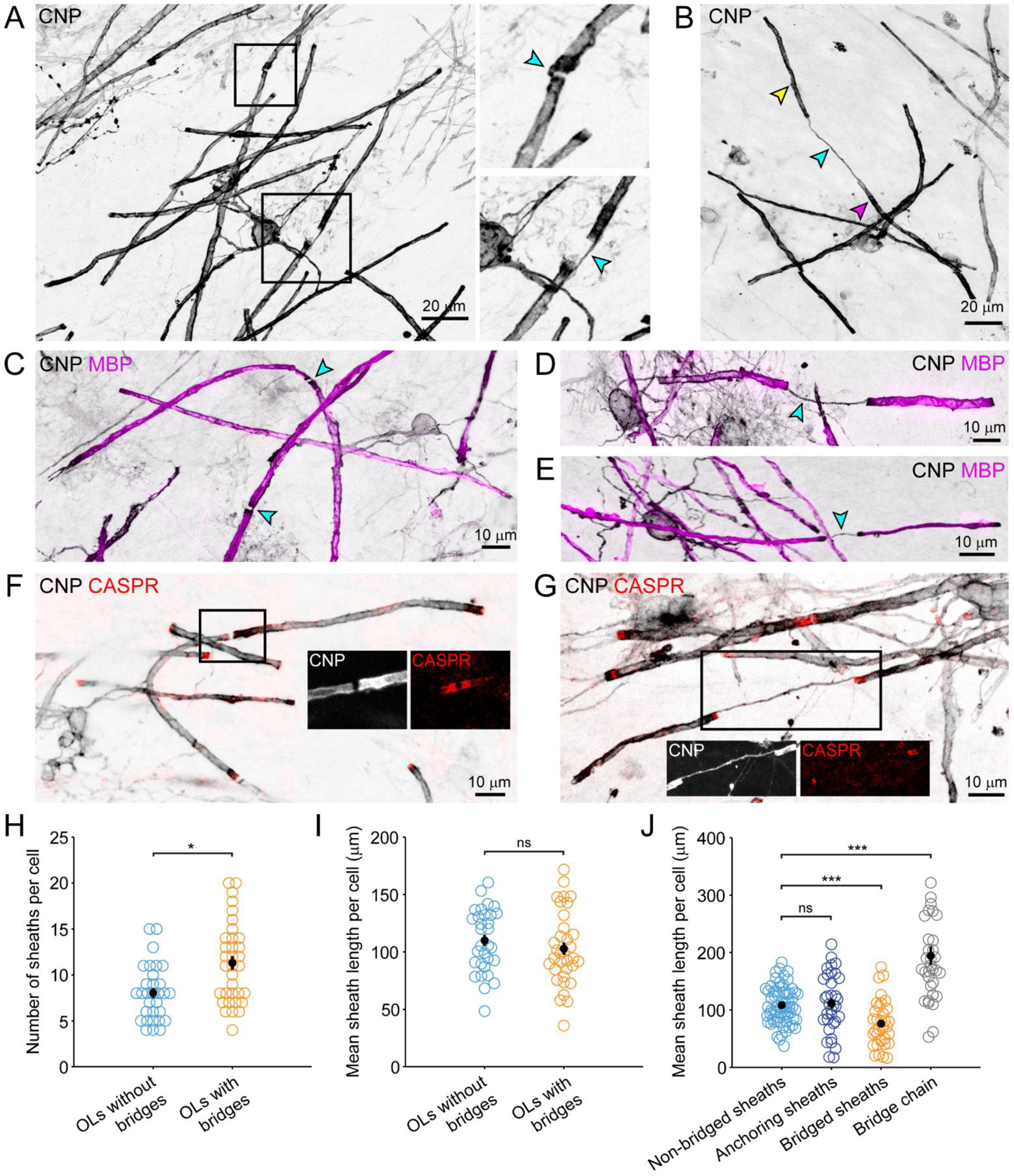
Oligodendrocytes within human organoids have paranodal bridges. (A) Representative image of single myelinating oligodendrocyte stained with CNPase. Squares highlight magnified paranodal bridge examples to the right. Arrowheads denote bridges.
(B) Example image of a pair of sheaths joined by a very long paranodal bridge. Arrowheads: magenta (bottom), anchoring sheath; cyan (middle), bridge; yellow (top), bridged sheath.
(C-E) MBP is excluded from bridged nodes of Ranvier.
(F-G) Representative images of paranodal bridges stained for CNPase (grayscale) and CASPR (red). Rectangles highlight region of single-channel insets.
(H) Quantification of the number of myelin sheaths per oligodendrocyte. Oligodendrocytes with bridged sheaths had a 31.2% increase in sheath number per cell compared to cells without bridges (95% confidence interval (CI): 9.9% to 56.7% p = 0.0027; GLMM with cell-line, conversion and organoid-ID included as random effects, n = 31 oligodendrocytes without bridges and n = 34 oligodendrocytes with bridges from 19 separate organoids across three hPSC lines).
(I) Mean sheath length per cell was similar between oligodendrocytes with and without paranodal bridges (Linear mixed effects regression with cell-line, conversion and organoid-ID included as random effects, n = 31 oligodendrocytes without bridges and n = 34 oligodendrocytes with bridges from 19 separate organoids across three hPSC lines).
(J) Quantification of average length of non-bridged sheaths (light blue), anchoring sheaths (dark blue), bridged sheaths (orange) and the total length of anchored and bridged sheaths (“bridged chain,” gray). No difference was found between non-bridged and anchored myelin sheaths. Bridged sheaths were 32% shorter than non-bridged sheaths (95% CI: 19% to 49% reduction; p < 0.001) and the total chain length was found to be 74% longer than canonical non-bridged myelin sheaths (95% CI: 63% to 93%; p < 0.001). Linear mixed effects regression with cell-line, conversion, organoid-ID and cell-ID included as random effects, n = 65 oligodendrocytes from 19 separate organoids across three hPSC lines).

While human myelinoid oligodendrocytes exhibit bona fide myelination, they do not replicate all aspects of the CNS environment. Therefore, to determine if paranodal bridges are also formed in the human brain, we performed CNP and CASPR immunostaining on postmortem human brain tissue from primary motor cortex (layer II/III). Remarkably, paranodal bridge-like structures between adjacent sheaths were frequently observed in these samples, with 32.6 ± 2% of all nodes exhibiting CNP immunoreactivity extending across NoR (118/251 NoR across all samples) (Supplemental Figure 4A,B). Together, these findings indicate that paranodal bridges are a highly conserved feature of oligodendrocyte myelin in zebrafish, mice, and humans.

### Paranodal bridges are modified paranodal loops continuous with the outer tongue of myelin

To determine how paranodal bridges connect myelin sheaths across NoR, we examined their ultrastructural features by serial electron microscopy (EM), using a high-resolution EM dataset of layer II/III from the young adult mouse visual cortex (38 days old) (Dorkenwald et al., 2019). For compacted myelin sheaths contained within the bounds of this volume, cytoplasmic extensions were frequently observed connecting paranodes across NoR (19/67 nodes) (Figure 6A-D, Supplemental Movie 1). In every instance, the paranodal bridge was continuous with the outer tongue of a mature myelin sheath on either side of the NoR. In addition, these bridges frequently formed a modified paranodal loop that extended down to the axonal membrane at the NoR (Figure 6A-D, yellow arrowheads), suggesting that paranodal bridges are formed by the first extensions of membrane that interact with axons, which later become the outermost layer of myelin. As observed through live imaging and immunohistochemistry, EM reconstructions confirmed that sheaths connected by bridges had no other cytoplasmic connection to an oligodendrocyte cell body.

**Figure 6.**
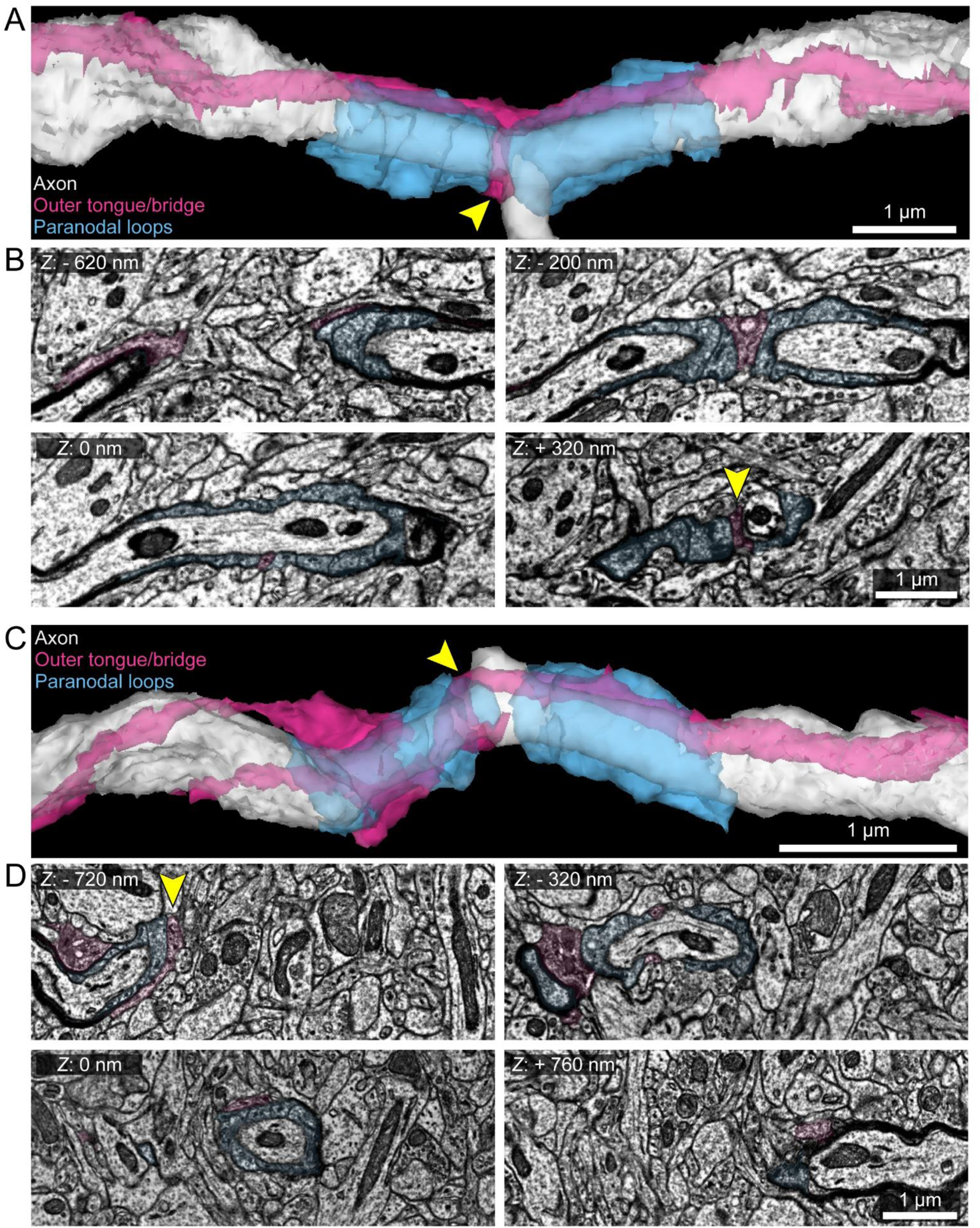
Paranodal bridges are continuous with the outer tongue and contact the axon as a modified paranodal loop. (A) 3D reconstruction of a paranodal bridge identified by electron microscopy (Dorkenwald et al., 2019) and reconstructed with Neuroglancer (https://neuroglancer-demo.appspot.com/). The cytoplasm of the paranodal loops are cyan, and the shared outer tongue which forms the paranodal bridge (band in between the two cyan structures, yellow arrowhead) is magenta. This paranodal bridge forms across a branch point in the axon.
(B) Individual frames making up the paranodal bridge reconstructed in *A*. The frame designated as z depth of 0 nm (bottom left frame) represents the center of the axon, and the other frames are from z planes above or below this frame. At 0 nm, the paranodal bridge is seen in cross section (magenta), indistinguishable from the other paranodal loops (cyan). At −200 nm and +320 nm (right frames), the paranodal bridge is observed on either side of the axon. Yellow arrowhead in bottom right frame represents approximate position of the same arrowhead in *A*. At −620 nm, the cytoplasm continuous with the magenta band at −200 nm can be observed passing over the other paranodal loops to the outside of both sheaths, which is continuous with the outer tongues of both sheaths (horizontal band running across *A*).
(C) Another 3D reconstruction as in *A*. In this example, the paranodal bridge is observed spiraling around the axon across the paranodal loops of the left sheath.
(D) As in *B*, at −320 nm the continuous magenta cytoplasmic channel is seen in cross section on either side of the axon, appearing as a paranodal loop. This cytoplasm is continuous with the magenta cytoplasm on the outer surface of the left sheath at −720 nm, and the right sheath at + 760 nm. Yellow arrowhead represents approximate position of the bridge as in *C*.

### Paranodal bridges occur frequently at axon branch points

Analysis of this serial EM dataset, in which the complex association of oligodendrocyte processes with axons could be visualized, revealed that the majority of paranodal bridges occurred at axon branch points (17/19 bridges) (Figure 6A, Supplemental Figure 5, Table 1), It is possible that the few paranodal bridges not associated with branch points are a remnant of earlier branching sites that were subsequently pruned with development (Buchanan et al., 2021; Portera-Cailliau et al., 2005; Yamahachi et al., 2009). Several examples of non-compacted, nascent sheaths that wrapped axons across branch points were observed in the volume (Supplemental Figure 6, Table 1). These results raise the possibility that growing sheaths extend past axon branches and subsequently split into two distinct sheaths during the compaction process, leaving the paranodal bridge to retain cytoplasmic continuity to the rest of the oligodendrocyte. We hypothesized this would most likely occur during the remodeling phase of myelination, which lasts for days after initial sheath elaboration has completed (Czopka et al., 2013; Orthmann-Murphy et al., 2020). Indeed, *in vivo* imaging of newly generated oligodendrocytes in the cortex of *Mobp-EGFP* mice revealed that some nascent sheaths (generated within the first 24-48 hr after the cell appeared) split to form two distinct sheaths over the course of several days, forming a bridged sheath and an anchoring sheath separated by a NoR (Supplemental Figure 7). Similar events were also observed over a timeframe of three days in zebrafish with mosaic expression of mbp:EGFP-CAAX (Supplemental Figure 8A); however, these bridges were often temporary and typically resolved into a continuous sheath within 24 hours (Supplemental Figure 8B). It is possible that the reduced probability of bridge stabilization reflects differences in the architecture of axons in the zebrafish spinal cord compared with those in the cortex (Auer et al., 2018; Koudelka et al., 2016), and that bridges may be more likely to form on neurons with highly arborized axons.

**Figure 7.**
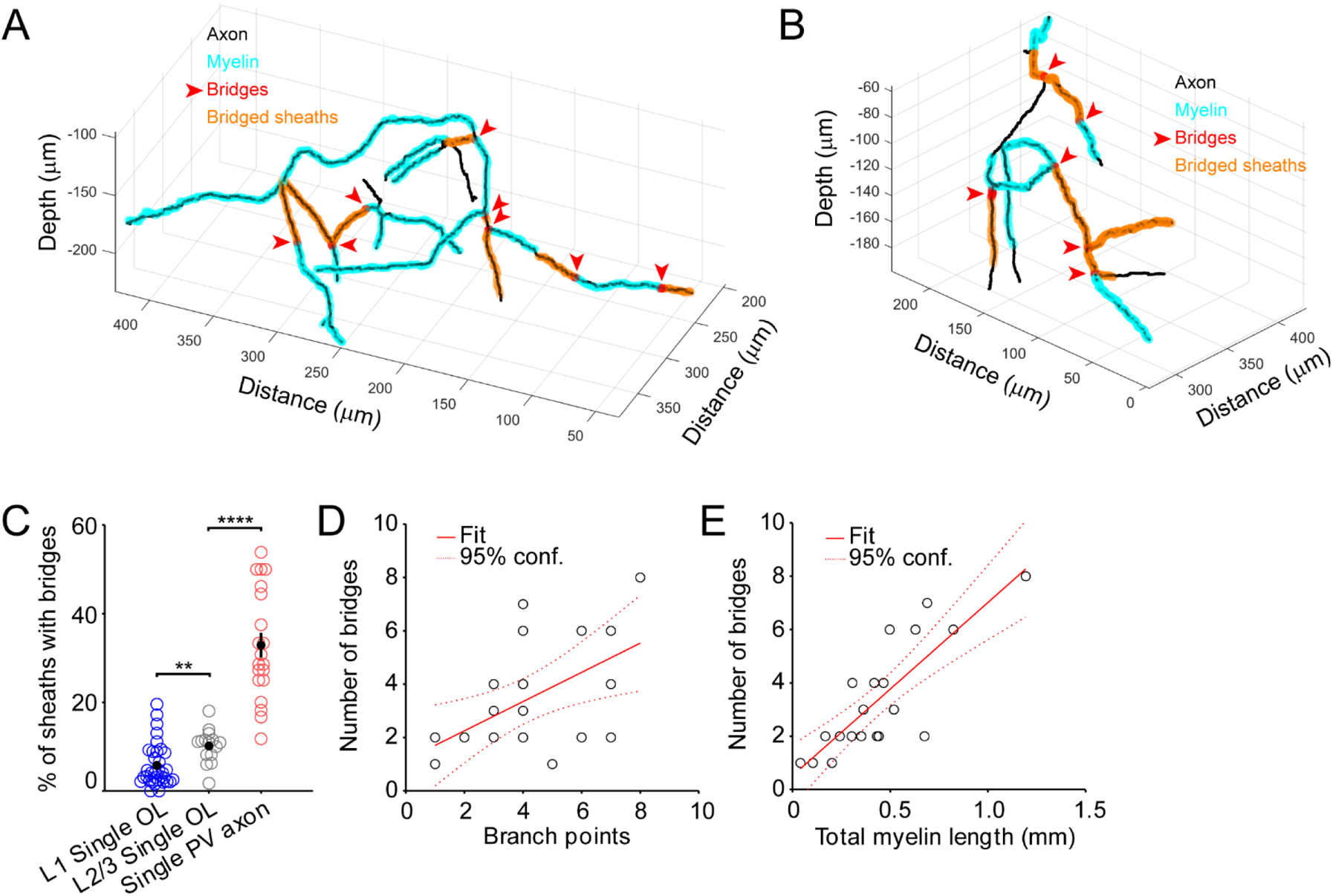
Paranodal bridges are overrepresented on PV axons and frequently span branch points. (A-B) 3D reconstructions of PV axons imaged *in vivo* in *PV-Cre; Ai9; Mobp-EGFP* somatosensory cortex. Depth is relative to pia.
(C) Individual PV axons have significantly higher proportions of bridged sheaths than are generated by individual oligodendrocytes (p = 1.2 × 10^−15^, two-sample two-tailed t-test).
(D) The number of paranodal bridges per PV axon positively correlates with the number of branch points of the axon (R^2^ = 0.31, p = 0.017).
(E) The number of paranodal bridges per PV axon positively correlates with the total myelin coverage of the axon (R^2^ = 0.67, p = 1.1 × 10^−5^).

**Table 1.**
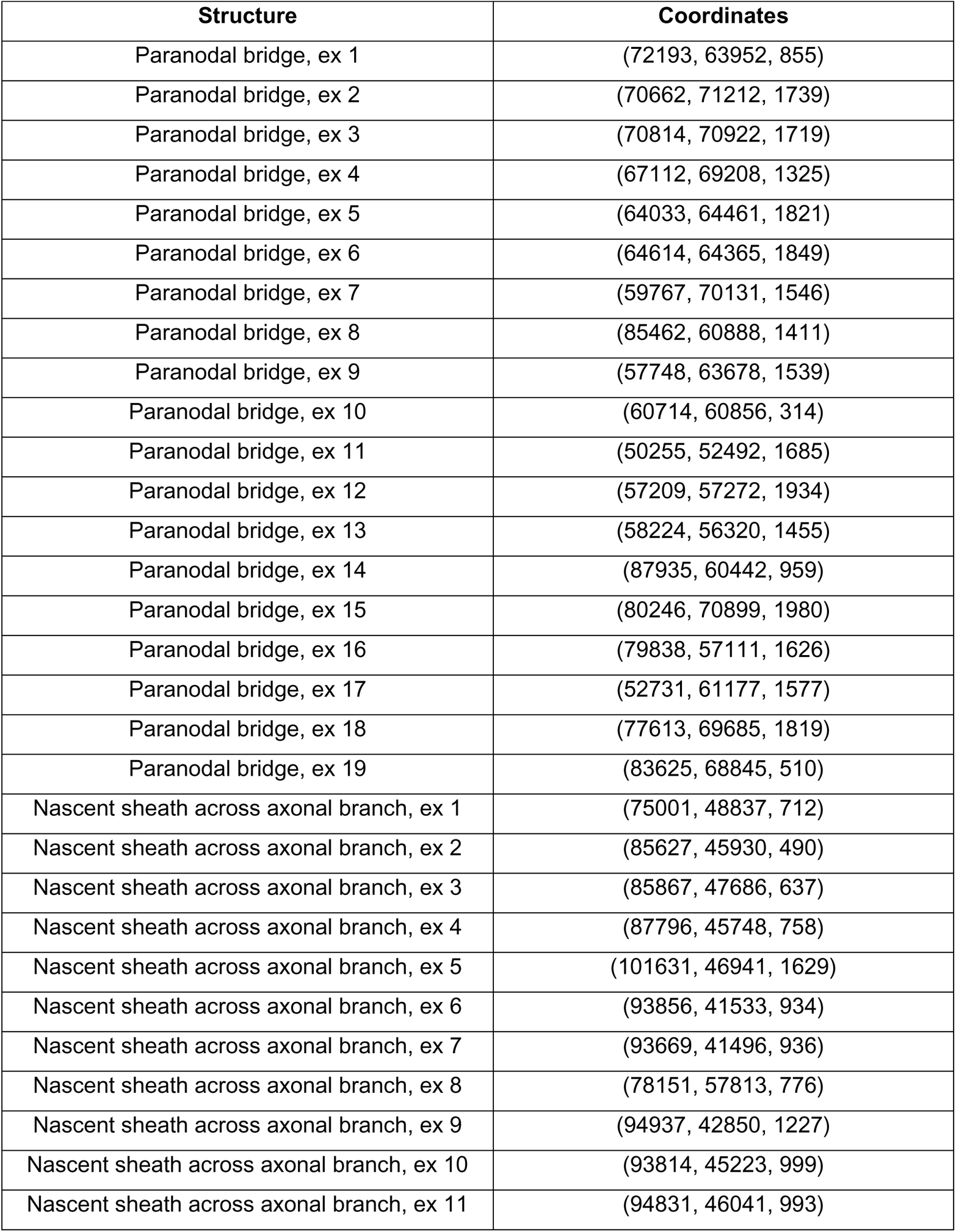
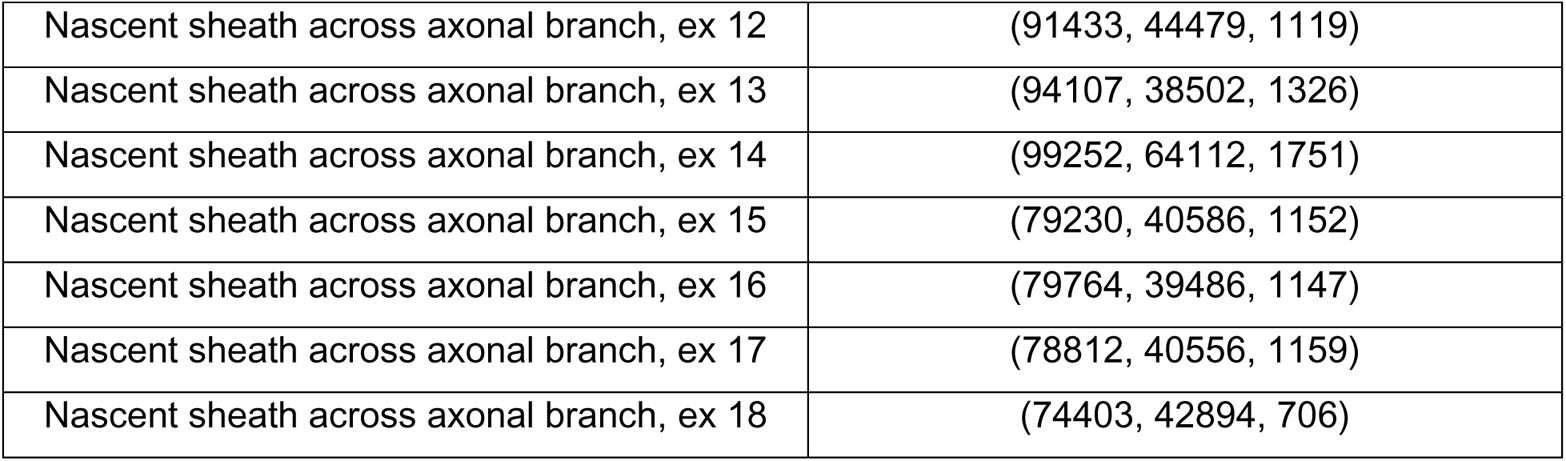
Coordinates of annotated structures in EM

**Table 2.**
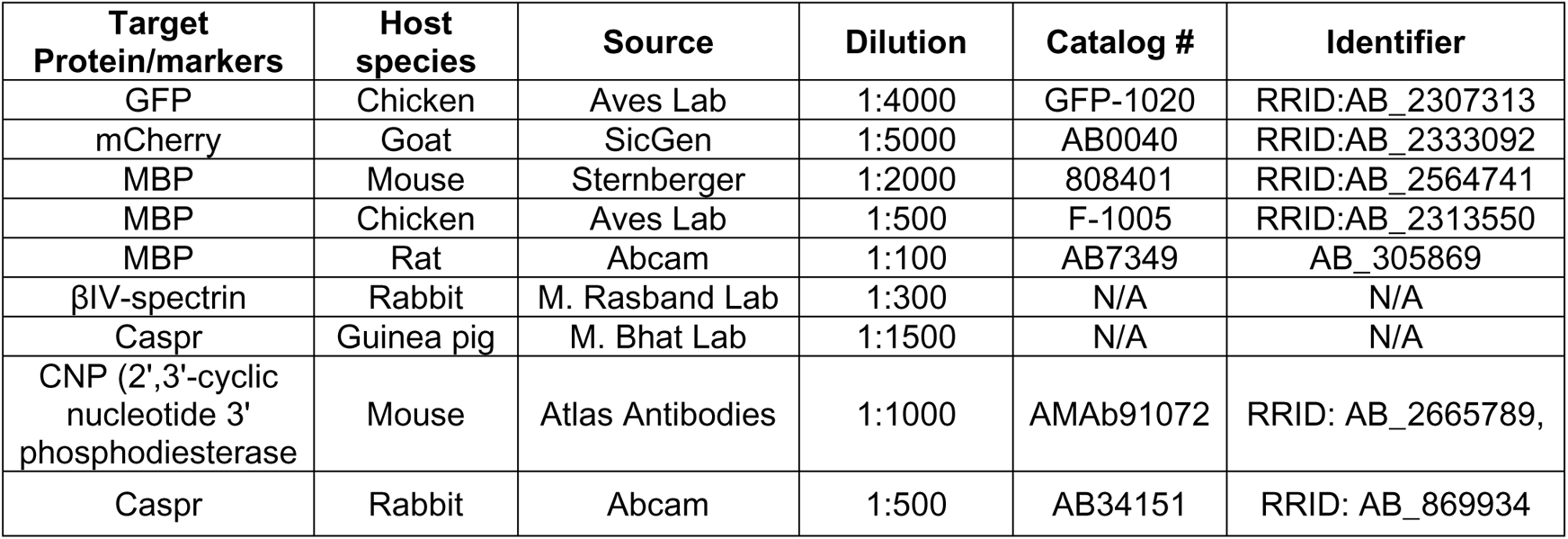
Primary antibodies

**Table 3.** Secondary antibodies

| Target Species | Conjugate | Source | Dilution | Catalog # | Identifier |
| --- | --- | --- | --- | --- | --- |
| Mouse | Cy5 | Jackson Immuno | 1:2000 | 715-175-151 | RRID:AB_2340820 |
| Goat | Cy3 | Jackson Immuno | 1:2000 | 705-166-147 | RRID:AB_2340413 |
| Chicken | Alexa 488 | Jackson Immuno | 1:2000 | 703-546-155 | RRID:AB_2340376 |
| Chicken | Cy5 | Jackson Immuno | 1:2000 | 703-006-155 | RRID:AB_2340347 |
| Rabbit | Cy3 | Jackson Immuno | 1:2000 | 711-165-152 | RRID:AB_2340817 |
| Rabbit | Alexa 647 | Jackson Immuno | 1:2000 | 711-605-152 | RRID:AB_2340820 |
| Guinea Pig | Cy3 | Jackson Immuno | 1:2000 | 706-165-148 | RRID:AB_2340461 |
| Guinea Pig | Alexa 647 | Jackson Immuno | 1:2000 | 706-605-148 | RRID:AB_2340477 |
| Rabbit | Alexa 488 | Thermo Fischer Scientific | 1:1000 | A-11008 | RRID: AB_143165 |
| Mouse | Alexa 568 | Thermo Fischer Scientific | 1:1000 | A-21144 | RRID: AB_2535780 |
| Rat | Alexa 488 | Thermo Fischer Scientific | 1:1000 | A-11006 | RRID: AB_2534074 |
| Rabbit | Alexa 647 | Thermo Fischer Scientific | 1:1000 | A-27040 | AB_2536101 |

**Table 5.** Software and Algorithms

| Name | Source | Identifier |
| --- | --- | --- |
| ZEN Blue/Black | Zeiss | RRID:SCR_013672 |
| Fiji | http://fiji.sc | RRID:SCR_002285 |
| Simple Neurite Tracer and SNT | https://imagej.net/SNT | RRID:SCR_016566 |
| Adobe Illustrator CS4 | Adobe | RRID:SCR_014198 |
| MATLAB | Mathworks | RRID:SCR_001622 |
| syGlass | IstoVisio | RRID:SCR_017961 |
| Prism | GraphPad | RRID:SCR_002798 |

### Paranodal bridges occur frequently on axons of PV interneurons

Given observations of the high incidence of paranodal bridges across axonal branch points, we wanted to explore the relative abundance of bridges along the highly branched PV interneurons of the cortex, which despite their short axonal lengths are among the most myelinated in the cerebral cortex (Call and Bergles, 2021; Micheva et al., 2016; Stedehouder et al., 2017; Zonouzi et al., 2019). Previous histological analysis using MBP immunolabeling revealed that the presence of myelin along PV interneuron axons is related to both the length and diameter of their discrete axon segments (Stedehouder et al., 2019), revealing that proximity to branch points negatively influences myelination. To visualize the organization of myelin along PV interneuron axons relative to branch points, we generated *Mobp-EGFP; PV-Cre; Ai9* mice in which oligodendrocytes and PV interneurons are cytoplasmically labeled with EGFP and tdTomato, respectively (Call and Bergles, 2021). *In vivo* two photon imaging within cortical layers I–II/III (∼60–230 µm below the pia) revealed that PV interneuron axons had an unusually large number of paranodal bridges (Figure 7A,B). Of 20 tdTomato+ axons traced from 9 mice, 30.1% of myelin sheaths were connected by paranodal bridge (68/222), with an average of 32.8 ± 3% bridged sheaths per axon (Figure 7C). This incidence of bridging was significantly higher than the overall proportion of bridges generated by oligodendrocytes within the same region of the cortex (9.6 ± 1%, p = 1.9 × 10^−7^, two-sample two-tailed t-test with Bonferroni correction) (Figure 7C), which myelinate axons of several different neuron subtypes (Call and Bergles, 2021; Micheva et al., 2016; Zonouzi et al., 2019), supporting the conclusion that the incidence of paranodal bridges on highly myelinated PV interneurons is influenced by their extensive axon branching.

Oligodendrocytes within layer II/III, where there is a high density of PV axons, exhibited a higher rate of bridged sheath formation than oligodendrocytes in layer I (p = 0.007, unpaired two-sample t-test with Bonferroni correction) (Figure 7C). Layer I PV interneuron axons are myelinated at a rate several times lower than in layer II/III (Micheva et al., 2016) and Layer I myelinated axons branch very infrequently (Call and Bergles, 2021), suggesting that regional differences in axonal morphology influence paranodal bridge formation. Indeed, 65 ± 8% of bridged sheaths from individual layer II/III oligodendrocytes were on highly branched PV interneuron axons. Along PV interneuron axons, 69.1% of paranodal bridges crossed axon branch points, and the number of bridges per axon was positively correlated with both the extent of axon branching (R^2^ = 0.31; Figure 7D) and total myelin coverage (R^2^ = 0.67; Figure 7E). This analysis likely underestimates the incidence of branching, due to resolution limits and the challenge of observing thin axons within highly dense neuropil *in vivo*. The enrichment of paranodal bridges on PV interneurons suggests that their formation is determined by the characteristics of these neurons, rather than the features of a distinct subset of oligodendrocytes.

### Bridged sheaths are more susceptible to degeneration in the aged brain

Normal aging is associated with a loss of myelin in humans, and oligodendrocytes have been shown to lose myelin sheaths and degenerate in older mice (Hill et al., 2018; Wang et al., 2020). To determine whether bridged sheaths exhibit greater vulnerability within the aged CNS, we used *in vivo* imaging to examine the dynamics of myelin sheaths over four to nine weeks in the somatosensory cortex of aged *Mobp-EGFP* mice (P585-P594) (Figure 8A-B). Sheaths degenerated at low rates in all regions examined (average of 8 ± 1 sheaths lost per 212 µm × 212 µm × 100 µm volume, range 1-18, n = 14 volumes, 3 mice) (Figure 8A,B).

**Figure 8.**
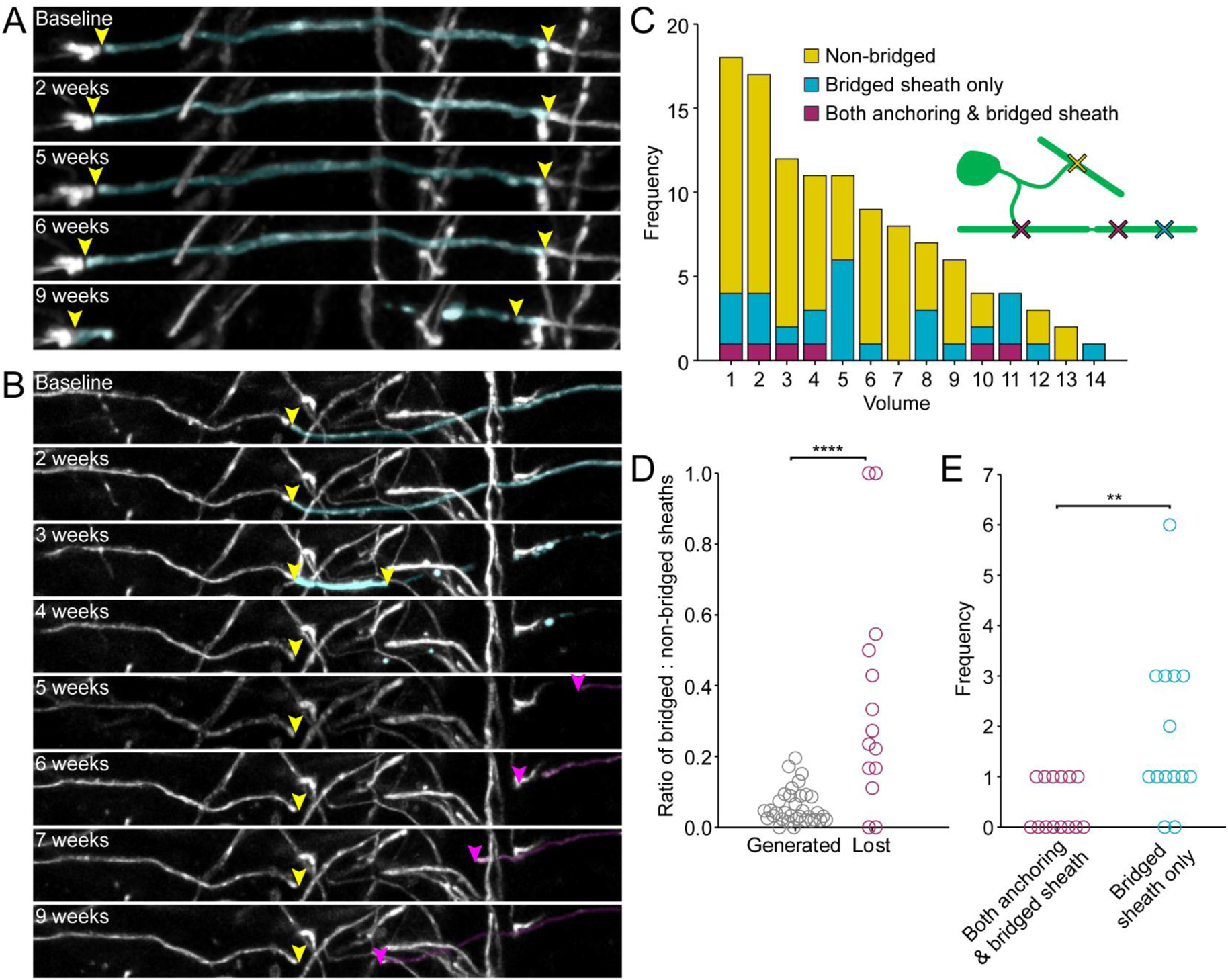
Sheaths connected by paranodal bridge are vulnerable to degeneration in old age. (A) Example of a bridged sheath (highlighted cyan) that degenerates in the aged *Mobp-EGFP* mouse cortex over the course of nine weeks of imaging. Yellow arrowheads mark nodes of Ranvier. Note that there is no cytoplasmic process connected to the highlighted sheath.
(B) Another example of a degenerating bridged sheath (cyan). Note that at three weeks, the right part of the sheath has degenerated, but the left part still connected by bridge to its neighboring sheath seems to partially stabilize before completely degenerating at four weeks. Following clearance of the bridged sheath, the neighbor on the right (magenta) overtakes the original position of the bridged sheath and has nearly reformed the left node by nine weeks. Arrowheads mark locations of paranodes.
(C) Histogram of the identity of degenerating sheaths in each quadrant. Inset schematizes each category (yellow, non-bridged; cyan, only the bridged sheath is lost; magenta, both the anchoring sheath and its bridged sheath are lost).
(D) Plot comparing the ratios of bridged:non-bridged sheaths of those generated in early life (pooled data from baseline and control cells from Figure 3B) and those lost in the aged brain. **** p = 2.1 × 10^−4^, Kruskal-Wallis.
(E) Frequency plot of data in C, only comparing sheaths within a bridged pair (cyan and magenta) ** p = 0.002, signed rank test.

Oligodendrogenesis has been reported in the aged cortex (Wang et al., 2020) and myelin sheath loss also occurs during initial maturation of oligodendrocytes (Orthmann-Murphy et al., 2020), raising the possibility that this sheath loss represents remodeling of newly generated oligodendrocytes. However, we did not observe oligodendrogenesis in these imaging volumes, indicating that these changes reflect the removal of internodes by existing, mature oligodendrocytes. Time lapse imaging revealed that bridged sheaths were almost 10 times more likely to degenerate than non-bridged sheaths (p = 1.87 × 10^−4^, two-sample Kolmogorov-Smirnov Test) (Figure 8C,D). Moreover, when comparing the relative degeneration of pairs of sheaths (i.e. loss of bridge chain versus loss of bridged sheath only), bridged sheath-only degeneration was much more frequent (bridge chain loss: 0.4 ± 0.1 sheaths per volume, range: 0–1; bridged sheath only loss: 2 ± 0.4 sheaths per volume, range: 0–6, p = 0.0020, paired signed rank test) (Figure 8E). Together, these data indicate that distal sheaths connected by paranodal bridges are more susceptible to degeneration in the aged brain.

## DISCUSSION

Myelin was among the first structures in the nervous system to be identified (Boullerne, 2016; van Leeuwenhoek, 1719). Its cellular origin was defined when individual oligodendrocytes were visualized using the Golgi stain (del Rio Hortega, 1922; Penfield, 1924), and its remarkable ultrastructure of concentric membrane wraps described though X-ray diffraction and electron microscopy shortly thereafter (Geren, 1954; Schmitt et al., 1935). Although the study of myelin has spanned many decades, this analysis has recently been reinvigorated by the ability to fluorescently label oligodendrocytes, myelin components, and recipient axons, allowing the dynamics of myelination to be described with high temporal and spatial resolution *in vivo*, the pattern of myelin along axons of distinct neuron subtypes to be defined, and the process of oligodendrocyte and myelin regeneration to be analyzed longitudinally within the intact nervous system. Using these approaches in multiple model systems, we discovered an unexpected structural feature of myelin in the CNS, in which individual myelin sheaths extend thin cytoplasmic processes across NoR, which we term paranodal bridges, to establish concatenated sheaths along a single axon, and in some cases bridge sheaths between nearby axons. This mechanism of linking internodes is highly conserved across vertebrates and enables myelin to cross axonal branch points, increasing the extent of myelin coverage along individual axons without additional oligodendrogenesis.

### Myelin addition through paranodal bridges

Myelination of axons in the CNS proceeds in a largely opportunistic manner, in which differentiating OPCs produce highly branched processes that interact with nearby axons, extending membrane sheaths along and around axon segments that exhibit appropriate features. Morphological reconstructions suggest that each sheath is produced by a (often branched) cytoplasmic process of an oligodendrocyte in a one-to-one ratio, with cytoplasmic connections to the soma emanating at varying locations along each internode (Butt and Ransom, 1993; Czopka et al., 2013; Murtie et al., 2007; Orthmann-Murphy et al., 2020). Several studies have reported atypical cytoplasmic processes or filopodia extending from oligodendrocytes at the paranodes of developing myelin sheaths (Haber et al., 2009; Hardy and Friedrich, 1996; Ioannidou et al., 2012; Toth et al., 2021); however, these structures were thought to represent transient extensions, pruned through the remodeling process that occurs during early stages of myelination (Czopka et al., 2013; Haber et al., 2009; Orthmann-Murphy et al., 2020). Our studies suggest that these structures may allow oligodendrocytes to extend processes around axon branch points and even jump to other nearby axons. Once firmly established, the two myelin segments joined by paranodal bridges remain extremely stable, persisting for > 2 months in the mouse brain, consistent with the limited turnover of myelin in the adult CNS (Hill et al., 2018; Hughes et al., 2018; Tripathi et al., 2017; Yeung et al., 2014; Young et al., 2013), indicating that these concatenated sheaths are not transient developmental structures, but rather a stable means to extend myelin along individual axons.

Although the proportion of sheaths connected by paranodal bridges (“bridged sheaths”) constitute less than 10% of total sheath production by most oligodendrocytes, the consistent production of bridged sheaths by oligodendrocytes yields considerable additional myelin within the cerebral cortex. For example, a 100 µm cube of layer I of the mouse somatosensory cortex contains approximately 150 sheaths (Orthmann-Murphy et al., 2020), each with a length of ∼70 µm. If this is extrapolated to the entire layer I of the barrel field (approximately 100, 100 µm cubes), there is approximately 1,050 mm of myelin, of which 70 mm would be formed using paranodal bridges. As the average cortical oligodendrocyte produces about 3 mm total myelin by length (Orthmann-Murphy et al., 2020) (Figure 3C), the formation of bridged sheaths represents the output of ∼23 oligodendrocytes within layer I of the barrel field, substantially reducing the number of oligodendrocytes required to establish this pattern of myelin. Given that the rate of bridged sheath formation is increased in deeper layers with higher rates of axon branching, the total bridged myelin content within the entire mouse cortex may be much greater, and even higher in the human cortex where the rate of bridging was ∼3 times that observed in mice.

Oligodendrocytes have only a limited range of myelinogenic potential and even under optimal circumstances, such as during development or after extensive demyelination where there is abundant axonal territory to myelinate, enhanced myelin generation by individual oligodendrocytes is minimal (Orthmann-Murphy et al., 2020). The limited production of myelin by each oligodendrocyte, the low rates of oligodendrocyte generation, and the presence of highly branched axons that constrain sheath formation (Stedehouder et al., 2019), create major challenges for myelinating even highly permissible axons in the cortex. If oligodendrocytes were only able to form a single internode from each cytoplasmic process, processes would have to branch further to add additional sheaths or additional oligodendrocytes would have to be generated to produce enough sheaths (of shorter lengths) to achieve continuous myelination across all branches. Moreover, in sparsely myelinated regions where individual oligodendrocytes can be separated by tens to hundreds of micrometers, additional oligodendrocytes would have to be formed to generate the single sheath needed to span the gap between cells (Supplemental Figure 2). As encounters between axons and the processes of premyelinating oligodendrocytes appear to be stochastic, a means would also have to exist to guide additional processes to these neighboring portions of axons.

Myelination provides significant savings in energy required to propagate action potentials, by replacing the need for continuous regeneration of action potentials with passive propagation and infrequent regeneration during saltatory conduction. However, oligodendrogenesis and myelination are energetically expensive, requiring extensive lipid and protein synthesis in a short period of time. It has been estimated that the initial energy cost required to invest in myelination is equivalent to several months of neuronal activity without myelination (Harris and Attwell, 2012). Thus, in the cortex where myelination is often sparse and discontinuous and only a small subset of axons is extensively myelinated, it may have been less advantageous to extend myelin by producing additional oligodendrocytes. The evolution of sheath production through paranodal bridges may have provided substantial advantages in energy savings, and as a result was conserved across vertebrate evolution.

### Paranodal bridges enable myelin continuity across axonal branch points

Volumetric EM analysis of the developing visual cortex revealed that nascent sheaths frequently extended past axonal branch points (Supplemental Figure 6), and mature paranodal bridges were remarkably frequent at axonal branch points of PV interneurons *in vivo* (Figure 7). In cortical layer II/III where the rate of PV interneuron myelination is highest and oligodendrocyte density is lowest, paranodal bridges were observed on PV interneuron axons at rates ∼3-fold higher than they are generated by individual oligodendrocytes, each of which form myelin on axons from different neuron subtypes (Call and Bergles, 2021; Micheva et al., 2016; Stedehouder et al., 2017; Zonouzi et al., 2019). Given the high propensity for bridges to be found spanning axon branch points, paranodal bridges present along some unbranched axons may represent sites where branches once emerged but were later pruned away (Buchanan et al., 2021; Portera-Cailliau et al., 2005; Yamahachi et al., 2009).

Myelination not only enhances the speed of action potential propagation, but can also enhance the fidelity of action potential propagation by preventing failures at branch points, which have particularly low safety factors for continued propagation (Manor et al., 1991; Parnas and Segev, 1979). Beyond the extreme case of conduction block, branch points have the potential to create delays in conduction in the daughter branches (Grossman et al., 1979a, 1979b). For axons that branch dozens of times, control of action potential arrival at all postsynaptic targets could be easily lost if conduction fidelity and relative delays between branch points were not tightly regulated. Even small delays can have profound impacts on oscillation synchrony (Pajevic et al., 2014). Thus, myelinating across branch points using paranodal bridges may represent an efficient way to preserve conduction with limited resources.

### Evolutionary conservation of myelin sheath concatenation

Concatenation of multiple myelin sheaths by a paranodal bridge was a consistent feature of oligodendrocytes in all three vertebrate systems examined in this study. While bridged sheaths in the adult mouse cortex were highly stable once formed, zebrafish paranodal bridges were dynamic, and were observed to form and dismantle over the course of several days. Additionally, in both fish and human myelinoids, many oligodendrocytes did not form paranodal bridges, while nearly every mouse oligodendrocyte exhibited bridged sheaths. These differences could be due to the relatively immature developmental environment or spinal cord origin of oligodendrocytes in larval zebrafish and myelinoid oligodendrocytes relative to the mouse cortex, and/ or differences in axonal structure (e.g. branching) within these tissues. Nonetheless, this comparative analysis highlights that extension of myelin through paranodal bridges is stochastic, raising the possibility that it may be modified by life experience.

### Aging dependent vulnerability of bridged myelin sheaths

During myelin compaction, cytoplasm is extruded from the myelin lamella, a process that is critical for establishing the insulating properties of myelin (Snaidero et al., 2017, 2014). The remaining few cytoplasmic channels are used for continued transport of materials necessary for sheath maintenance. However, it has been hypothesized that reparative processes could be impaired by this structural bottleneck when faced with metabolic stress or myelin damage (Hagemeyer et al., 2012; Lappe-Siefke et al., 2003; Saab and Nave, 2017), contributing to enhanced vulnerability of oligodendrocytes to metabolic insults. Bridged sheaths are connected to the rest of the oligodendrocyte exclusively through the paranodal bridge, a thin extension of the myelinic channel through which components necessary to sustain the distal sheath need to travel. Bridged sheaths tended to exist at the periphery of an oligodendrocyte’s territory (Supplemental Figure 1), increasing the distance required to deliver proteins, mRNA and metabolites. Transport defects in long axons have been described in many neurodegenerative diseases, including Alzheimer’s disease and multiple sclerosis (Coleman, 2005). Similarly, paranodal bridges may place constraints on homeostatic mechanisms, such as protein and organelle trafficking, that experience greater dysfunction in the aging brain (Benveniste et al., 2019; Camandola and Mattson, 2017; Mattson and Arumugam, 2018; Uzor et al., 2020).

As a result of their enhanced vulnerability, bridged sheaths may contribute substantially to age-related demyelination, resulting in gaps along otherwise continuously myelinated axons and alter conduction across axon branch points. Such a scenario could be particularly detrimental to cortical synchrony, as a substantial portion of myelin segments on PV interneurons are generated by paranodal bridges, and these cells are crucial for producing cortical oscillations associated with learning and memory retrieval (Cardin et al., 2009; Cobb et al., 1995; Klausberger et al., 2004; Ratnadurai-Giridharann et al., 2015; Wang and Buzsáki, 1996). In most cases, gaps left by bridged sheath degeneration would be expected to become persistent, as oligodendrogenesis is rare in the aged brain (Hill et al., 2018; Wang et al., 2020). It is possible that other CNS insults that impact metabolism and cellular maintenance, such as inflammation, hypoxia and increased production of free radicals (Aboul-Enein et al., 2003; Segovia et al., 2008; Ziabreva et al., 2010), may disproportionally affect bridged sheaths and contribute to cortical dysfunction in neurodegenerative diseases. Examination of the fate of bridged sheaths in these diverse contexts will reveal their contribution to overall myelin loss, inform how these changes alter cortical circuits, and ultimately help us develop strategies to reduce their vulnerability to promote myelin stability across the lifespan.

## METHODS

### Mouse care and use

All experiments involving mice were conducted in strict accordance with protocols approved the Animal Care and Use Committee at Johns Hopkins University, in compliance with federal regulations. Female and male adult *Mobp-EGFP* mice were used for experiments and randomly assigned to experimental groups. All mice were healthy and did not display any overt behavioral phenotypes. Mice were maintained on a 12-h light/dark cycle, food and water were provided ad libitum, and housed in groups no larger than 5. Mice were housed with at least one other cage mate when possible. Three *Mobp-EGFP* mice were aged to 1.5 years before being implanted with cranial windows for our aging experiment.

### Mouse cranial windows

Cranial window surgeries were performed as described previously (Orthmann-Murphy et al., 2020). Briefly, *Mobp-EGFP* mice were deeply anesthetized with isoflurane (5% at 1 L/min O_2_ induction, 1.5–2% at 0.5 L/min O_2_ maintenance) and their scalps shaved and cleaned. A portion of the scalp was removed and the underlying skull was cleaned and dried before cementing (Metabond) on a custom aluminum headplate. A 3-mm circle of skull was removed with a high-speed dental drill and replaced with a coverslip that was secured in place with VetBond and Krazy Glue. Mice recovered in their home cage on a heating pad and were monitored for at least an hour. Mice were imaged two to three weeks following window surgery.

### *In vivo* two-photon microscopy in mice

*In vivo* imaging was performed as described previously (Orthmann-Murphy et al., 2020). Briefly, *Mobp-EGFP* mice were deeply anesthetized under isoflurane (5% at 1 L/min O_2_) and then transferred to a custom stage on a Zeiss 710 microscope and clamped in place by their headplate where they remained under isoflurane (1.5–2% at 0.5 L/min O_2_ maintenance) for the remainder of the imaging session. Two-photon images were collected on a Zeiss LSM 710 or Zeiss LSM 880 microscope with a GaAsP detector and mode-locked Ti:sapphire laser (Coherent Ultra) tuned to 920 nm (*Mobp-EGFP* mice) or 1000 nm (*Mobp-EGFP; PV-Cre; Ai9* mice) with average power at the sample < 30 mW. A Zeiss coverslip-corrected 20X water-immersion objective (NA 1.0) was used to acquire 2048 × 2048 pixel (425 μm × 425 μm) z-stacks from the pia to depths of 110 μm or 230 μm (1-μm z step).

To quantify bridge frequency on PV axons, axons in layer II/III were fully traced using SNT in ImageJ from *in vivo* images of *Mobp-EGFP; PV-Cre; Ai9* mice. Myelin sheaths surrounding these axon traces were traced, including any paranodal bridges and cytoplasmic processes leading to oligodendrocyte cell bodies. Sheaths were labeled as either being anchoring, bridged, or undefined and bridges were labeled as either spanning a branch or not.

### Mouse cortical flatmount preparation

Flatmount preparation was performed as described previously (Call and Bergles, 2021). Briefly, deeply anesthetized mice (100 mg/kg w/w sodium pentobarbital) were transcardially perfused with 20–25 mL warm (30–35° C) PBS followed by 20–25 mL ice-cold 4% paraformaldehyde. Cortical mantles were dissected from the underlying brain structures, unrolled, placed between two glass slides separated by 1 mm, and postfixed in 4% paraformaldehyde at 4° C for 6–12 hours. Flattened cortices were removed from the clamped slides and stored in 30% sucrose in PBS for at least 24 hours until sectioned on a cryostat (Thermo Scientific Microm HM 550) at –20 ° C at thicknesses of 35–50 μm. Cryostat chucks were pre-frozen with TissueTek mounting medium and sectioned until flat. Flatmounts were removed from sucrose solution, covered with mounting medium dorsal-side down on a silanized glass slide, and frozen onto the prepared chuck. Care was taken to ensure complete horizontal sections were acquired by aligning the blade angle to the surface of the tissue.

Mice used for nodal component immunostaining were perfused only with 20–25 mL warm (30–35° C) PBS. Brains were dissected and lightly fixed for 30–60 minutes in 4% PFA. Flatmounts were lightly post-fixed in 4% PFA for 60 minutes in the clamped slide configuration. Flatmounts continued to be maintained in the clamped slide configuration during 30% sucrose incubation for at least 24 hours before sectioning.

### Mouse immunohistochemistry

Immunohistochemistry on mouse brain was performed on free-floating tissue sections preincubated in blocking solution (5% normal donkey serum, 0.3% Triton X-100 in PBS, pH 7.4) for up to 2 hours at room temperature, then incubated for 24–48 hours at 4° C or room temperature in primary antibodies. Sections were subsequently washed in PBS before being incubated in secondary antibodies at room temperature for 2–6 hours or overnight at 4° C. Sections were mounted on slides with Aqua Polymount (Polysciences). Specific antibodies used are listed in the Key Resources Table.

### Mouse image processing and analysis

Image registration, processing, and tracing of oligodendrocyte morphologies was performed as described previously (Orthmann-Murphy et al., 2020). High resolution imaging of individual bridged nodes was performed on a Zeiss 800 or 880 in Airyscan mode. Regions were ∼45 µm × 45 µm in xy with a resolution of ∼1800 × 1800 pixels. Maybe more context here for nature/ origin of tissue. Z stacks ranged in depth, but had z steps of 0.18 µm. For analysis of degenerating sheaths in aged mice, individual regions acquired with a 20x objective were subdivided into quadrants of 212 µm × 212 µm × 100 µm volumes prior to beginning analysis. Quadrants that had overlying blood vessels, bone, or thickened meninges during the course of imaging were excluded from analysis. Loss of individual sheaths was detected in syGlass volumetrically by observing 10-20 µm-thick slices at a time and continuously rotating through timepoints. Lost sheaths were verified in ImageJ and their identities (bridged or non-bridged) were then determined. The density of myelination and abundance of lipofuscin at these ages were substantial, preventing accurate tracing of full morphologies of individual oligodendrocytes.

However, we were able to distinguish bridged sheaths by their lack of intersecting cytoplasmic process between paranodes. Neighboring sheaths that degenerated simultaneously were considered the anchoring sheath of the bridged pair.

### Electron microscopy analysis of mouse cortex

We used the publicly available 250 µm × 140 µm × 90 µm EM volumetric dataset of a P36 mouse visual cortex layer II/III, acquired at a resolution of 3.58 nm × 3.58 nm × 40 nm (Dorkenwald et al., 2019). This dataset was automatically segmented with machine learning, and neuronal structures were subsequently validated manually through the Machine Intelligence from Cortical Networks (MICrONS) program (https://microns-explorer.org/). It was annotated with the online Neuroglancer interface (https://github.com/google/neuroglancer). Segmentation is based on cytoplasmic connectivity, and thus myelin membrane is not itself segmented, and segmentations of cytoplasmic channels within the sheath are fragmented. Thus, annotation consisted of finding nodes at low resolution by eye, and then activating segmented meshes associated with the paranodal loops and connecting these segmentations manually with adjacent cytoplasmic channels. In most cases, segmentation of paranodal bridges was continuous across both sheaths, but these were verified manually by tracing the cytoplasm between frames and confirming continuity.

Not all myelin sheaths were fully contained within the bounds of the volume. When possible, lack of a direct cytoplasmic process of one of the sheaths connected by paranodal bridge was confirmed by following the entirety of the outer tongue between paranodes of each sheath. Paranodal bridges identified in this dataset may be found at the coordinates listed in Table 1. These labels are the same as those used in Supplemental Figure 5.

### Zebrafish care and use

All zebrafish studies were carried out with approval from the UK Home Office according to their regulations under the following project licences: 70/8436 and PP5258250. Zebrafish were maintained in the Queen’s Medical Research Institute BVS Aquatics Facility at the University of Edinburgh. Adult zebrafish were maintained by aquatics staff under standard conditions on a 14 hours light, 10 hours dark cycle. Zebrafish embryos were maintained at 28.5°C in 10 mM HEPES buffered E3 embryo medium or in conditioned aquarium water with methylene blue. Larval zebrafish were analysed between 4-7 dpf, before zebrafish undergo sexual differentiation.

### *In vivo* confocal microscopy in zebrafish

To fluorescently label the myelin sheaths of single oligodendrocytes fertilised zebrafish eggs were injected at the single cell stage with 1 nl of 10 ng/µl pTol2-mbp:EGFP-CAAX plasmid DNA (Czopka et al., 2013) and 50 ng/µl Tol2 transposase mRNA. Zebrafish were screened to identify isolated fluorescently labelled oligodendrocytes from 3 dpf. To screen for fluorescently labelled oligodendrocytes, larval zebrafish were first anesthetised with MS222 before mounting them in 1.5% low melting point agarose on glass coverslips. Once zebrafish were anaesthetised and mounted, oligodendrocytes in the spinal cord were selected for imaging. Z-stacks of oligodendrocyte were acquired using the LSM880 confocal microscope with Airyscanner fast and a 20x objective (Zeiss Plan-Apochromat, NA = 0.8). Z-stacks were acquired with an optimal z-step for each experiment.

### Zebrafish image analysis

To quantify the number and lengths of myelin sheaths and paranodal bridges the segmented line tracing tool in Fiji Image J was used. Oligodendrocytes were analysed throughout the depth of each z-stack per cell. No cells were excluded from analyses unless there was too much myelin overlapping from neighbouring cells to reliably quantify myelin sheaths and paranodal bridges. One oligodendrocyte was analysed per zebrafish unless otherwise specified in figure legends.

### Human myelinoid generation and processing

The human pluripotent stem cell-lines used in this study were obtained with full Ethical/Institutional Review Board approval by the University of Edinburgh and validated using standard methods including chromosomal analysis, pluripotency and absence of plasmid integration. The iPSC lines CS02iCTR-NTn1 (male) and CS25iCTRL-18n2 (male) were obtained from Cedars-Sinai and the embryonic stem cell-line SHEF4 (male) was obtained from the UK Stem Cell Bank. The maintenance of human pluripotent stem cells and generation of myelinoid cultures has been described recently (James et al., 2021). Briefly, cells were maintained in Essential 8 medium before being lifted into suspension and patterned towards the pMN domain of the developing spinal cord. Spheroids containing ventral, caudal neuroepithelial cells were then patterned towards a glial cell-fate using PDGF-AA before being transferred onto PTFE-coated Millicell Cell Culture Inserts (Merck) and maintained until cultures were 19 weeks old (corresponding to MI-12 in James et al., 2021).

Myelinoids were fixed in 4% PFA, washed, then permeabilized in 0.25% triton-X-100 in PBS for 40 minutes and blocked in 10 % normal goat serum (Vector Laboratories) + 0.25% triton-X-100 for 2 hours at RT. For CNP immuno-staining, myelinoids were incubated in citrate buffer (pH 6) at 95° C for 20 minutes followed by a further hour in blocking solution. Primary antibodies rat anti-MBP, mouse anti-CNP and rabbit anti-CASPR were incubated overnight at 4° C in blocking solution. After washing in PBS (3x 20 min), secondary antibodies (goat anti-rat, goat anti-mIgG2b and goat anti-rabbit) were incubated for 2 hours at room temperature in blocking solution. Myelinoids were stained with DAPI, washed in PBS and whole-mounted onto microscope slides (Thermo Scientific) with FluorSave (Calbiochem) and No. 1.5 coverslips (Thermo Scientific). Images were captured using a Zeiss 710 confocal microscope and analysed in FIJI using the Cell Counter and Simple Neurite Tracer plugins for counting cells and tracing myelin sheath lengths, respectively.

### Human postmortem brain tissue

Post-mortem brain tissue (motor cortices) from people without neurological defects were provided by a UK prospective donor scheme with full ethical approval from the UK Multiple Sclerosis Society Tissue Bank (MREC/02/2/39) and from the MRC-Edinburgh Brain Bank (16/ES/0084). The clinical history was provided by R. Nicholas (Imperial College London) and Prof. Colin Smith (Centre for Clinical Brain Sciences, Centre for Comparative Pathology, Edinburgh). Table 6 provides the details of the samples that were used in the study. The mean age of the human tissue donors was 68.5 years. Tissue blocks of 2 cm × 2 cm × 1 cm were collected, fixed, dehydrated and embedded in paraffin blocks. 4-μm sequential sections were cut and stored at room temperature.

**Table 6.**
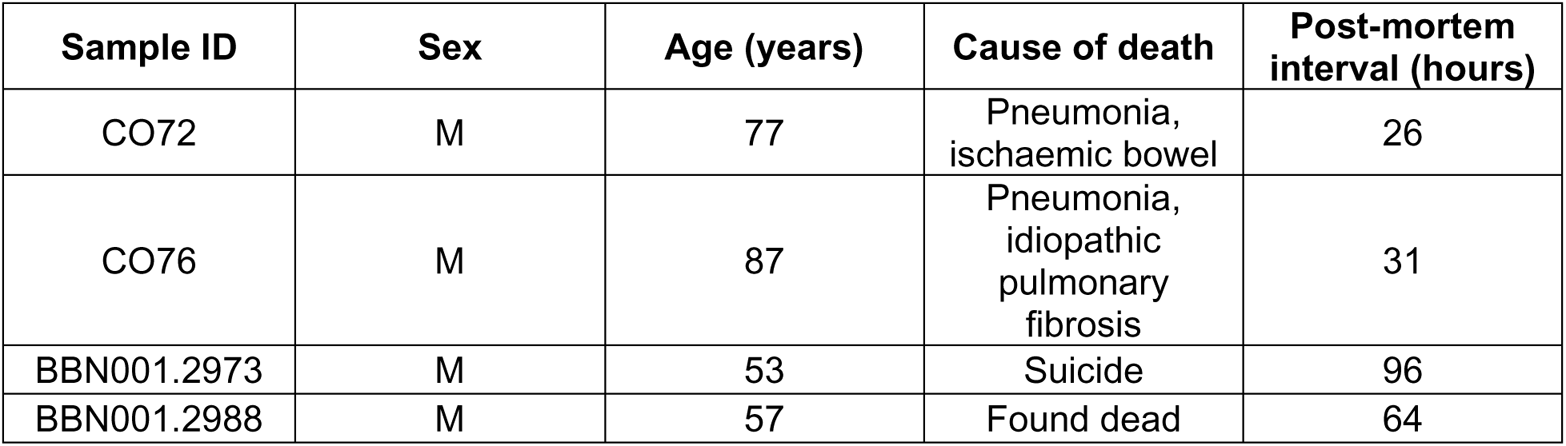
Details of human donor tissue

| Sample ID | Sex | Age (years) | Cause of death | Post-mortem interval (hours) |
| --- | --- | --- | --- | --- |
| CO72 | M | 77 | Pneumonia, ischaemic bowel | 26 |
| CO76 | M | 87 | Pneumonia, idiopathic pulmonary fibrosis | 31 |
| BBN001.2973 | M | 53 | Suicide | 96 |
| BBN001.2988 | M | 57 | Found dead | 64 |

### Immunohistochemistry on human tissue

Paraffin sections were rehydrated, washed in PBS and microwaved at high power for 15 minutes in Vector Unmasking Solution for antigen retrieval (H-3300, Vector Laboratories). The sections were then incubated with Autofluorescent Eliminator Reagent (2160, MERCK-Millipore) for 1 minute and briefly washed in 70% ethanol for 5 minutes. Image-iT® FX Signal Enhancer (I36933, Thermo Fisher Scientific) was subsequently applied for 30 minutes at room temperature, and then the sections were washed and blocked for 1 hour with 10% normal horse serum, 0.3% Triton-X in PBS. Primary antibodies were diluted in antibody diluent solution (003118, Thermo Fisher Scientific) and sections were incubated overnight at 4⁰ C in a humidified chamber. The next day the sections were incubated with Alexa Fluor secondary antibodies for 1 1/2 hours at room temperature, counterstained with Hoechst 33342 (62249, Thermo Fisher Scientific) for the visualization of the nuclei and mounted using Mowiol^®^ mounting medium (475904, MERCK-Millipore). The details of the antibodies used are listed in the Key Resources Table. Z-stack images were acquired from layers 2 and 3 of the human primary motor cortex with Leica TCS SP8 confocal microscope using a 63x objective. From each sample up to 14 different regions of ∼62 µm × 62 µm in xy with a resolution of ∼2048 × 2048 pixels and a system’s optimized z step were acquired and the average percentage of bridged NoR were quantified.

## DATA AVAILABILITY STATEMENT

All published image data, code, tools, and reagents will be shared on an unrestricted basis; requests should be directed to the corresponding authors. Raw tracing data files are available at https://github.com/clcall/Call_ParanodalBridge_2022 and summary data is included in the Source Data file. Source data are provided with this paper.

## CODE AVAILABILITY STATEMENT

MATLAB scripts and ImageJ macros are available at https://github.com/clcall/Call_ParanodalBridge_2022.

## SUPPLEMENTAL FIGURES

**Supplemental Figure 1.**
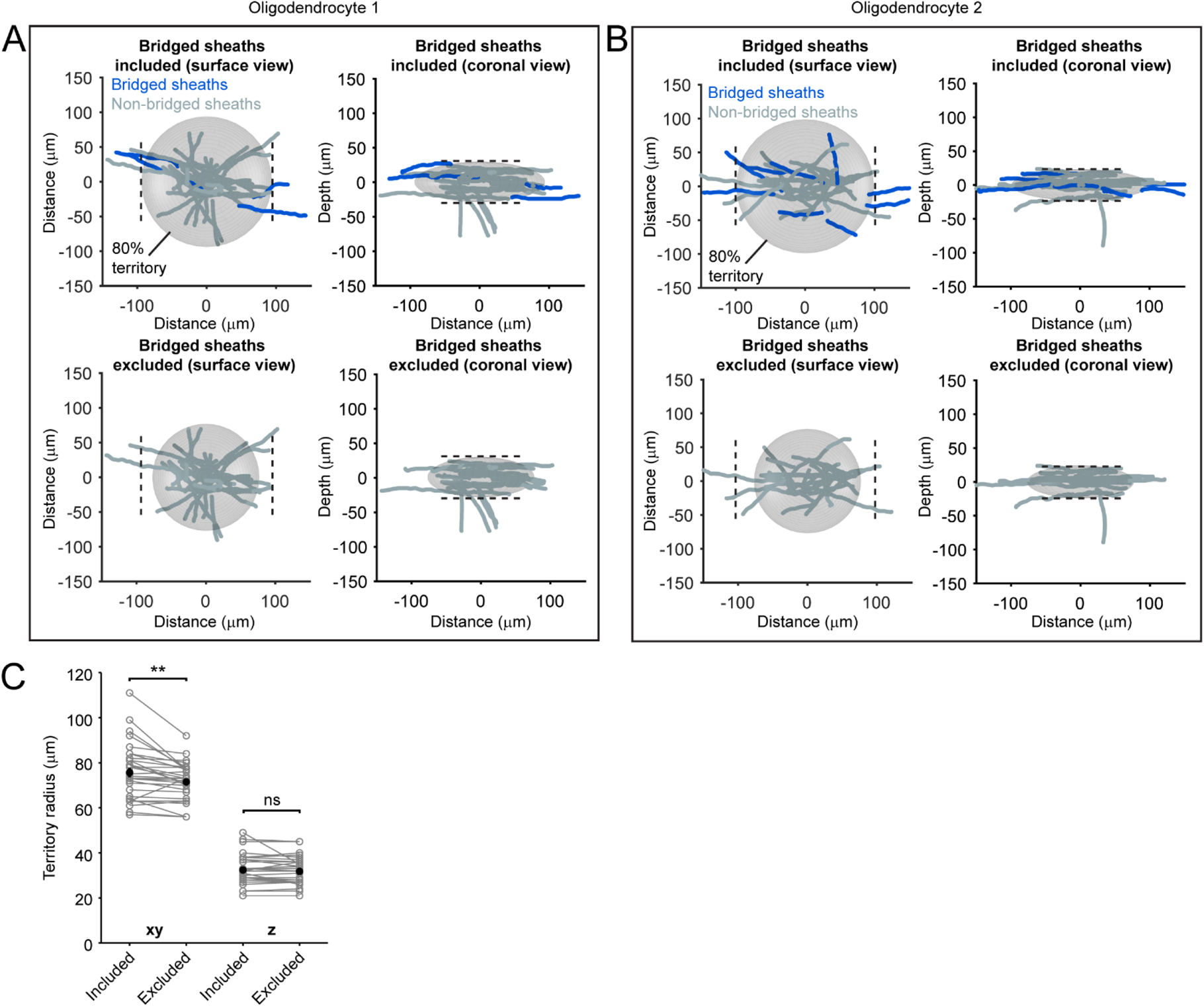
Bridged sheaths expand oligodendrocyte territory. (A-B) Sets of sheath cohort reconstructions from two oligodendrocytes. The top plots show all sheaths including bridged sheaths (highlighted dark blue) as viewed from the brain surface (left) and coronally (right). Best-fit territory ellipsoids encompassing ≥ 80% of total sheath content are superimposed. Bottom plots show only unbridged and anchoring sheaths of the same cell and the superimposed territory. Dotted lines indicate the diameter of territories with bridged sheaths included and are in the same relative positions in the top and bottom rows. (C) Quantification for best-fit territory radii in xy and z dimensions for all existing and control oligodendrocytes used in Figure 3. The xy (p = 0.002, paired two-tailed t-test with Bonferroni correction), but not z (p = 0.46, paired two-tailed t-test with Bonferroni correction) radii are significantly smaller when bridged sheaths are excluded.

**Supplemental Figure 2.**
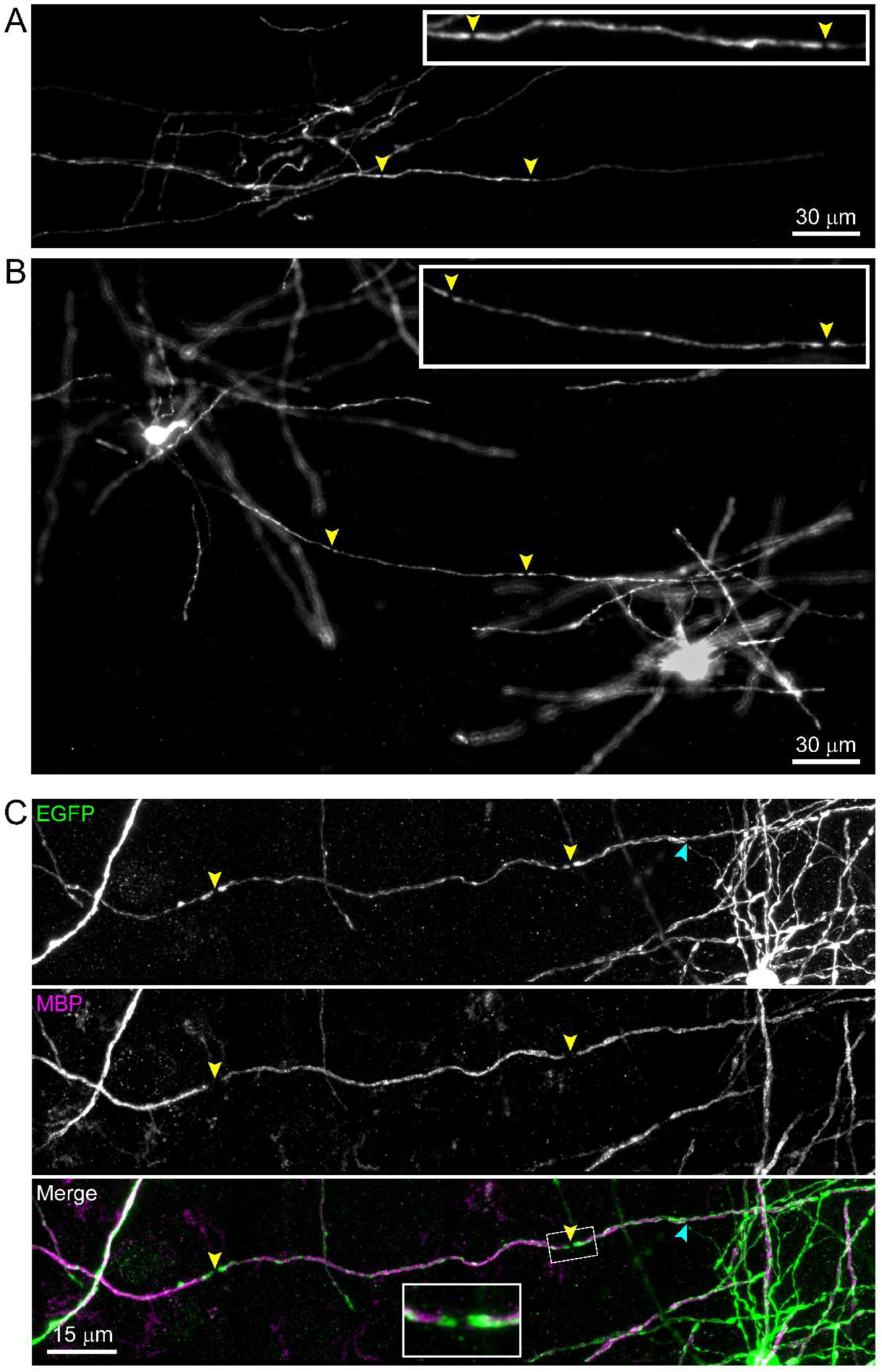
Paranodal bridges allow continuous myelination of select axons in sparsely myelinated regions. (A) Epifluorescence image of a partial cohort of sheaths from an oligodendrocyte (cell body not in section) in an *Mobp-EGFP* cortical flatmount in the temporal association area. A series of sheaths provide continuous myelination to one axon, and two paranodal bridges (yellow arrowheads) link three sequential sheaths from a single oligodendrocyte.
(B) As in *A*. This example shows two oligodendrocytes which share a single axon. The axon is continuously myelinated in between these cells, which would otherwise require a third oligodendrocyte if it were not for a bridged sheath provided by one of these cells.
(C) Similar to *A* and *B*, but a high resolution confocal z projection showing a continuously myelinated axon with one bridged sheath in the center. MBP immunostaining (center/magenta) shows two clear breaks along this axon, which are associated with nodes of Ranvier (yellow arrowheads), visible by the EGFP-positive doublets in the top panel. The inset in the bottom panel shows a magnified image of the node with a paranodal bridge. The blue arrowhead indicates the intersection between the cytoplasmic process and the anchoring sheath. ****p = 1.87E-4, two-sample Kolmogorov-Smirnov Test.

**Supplemental Figure 3:**
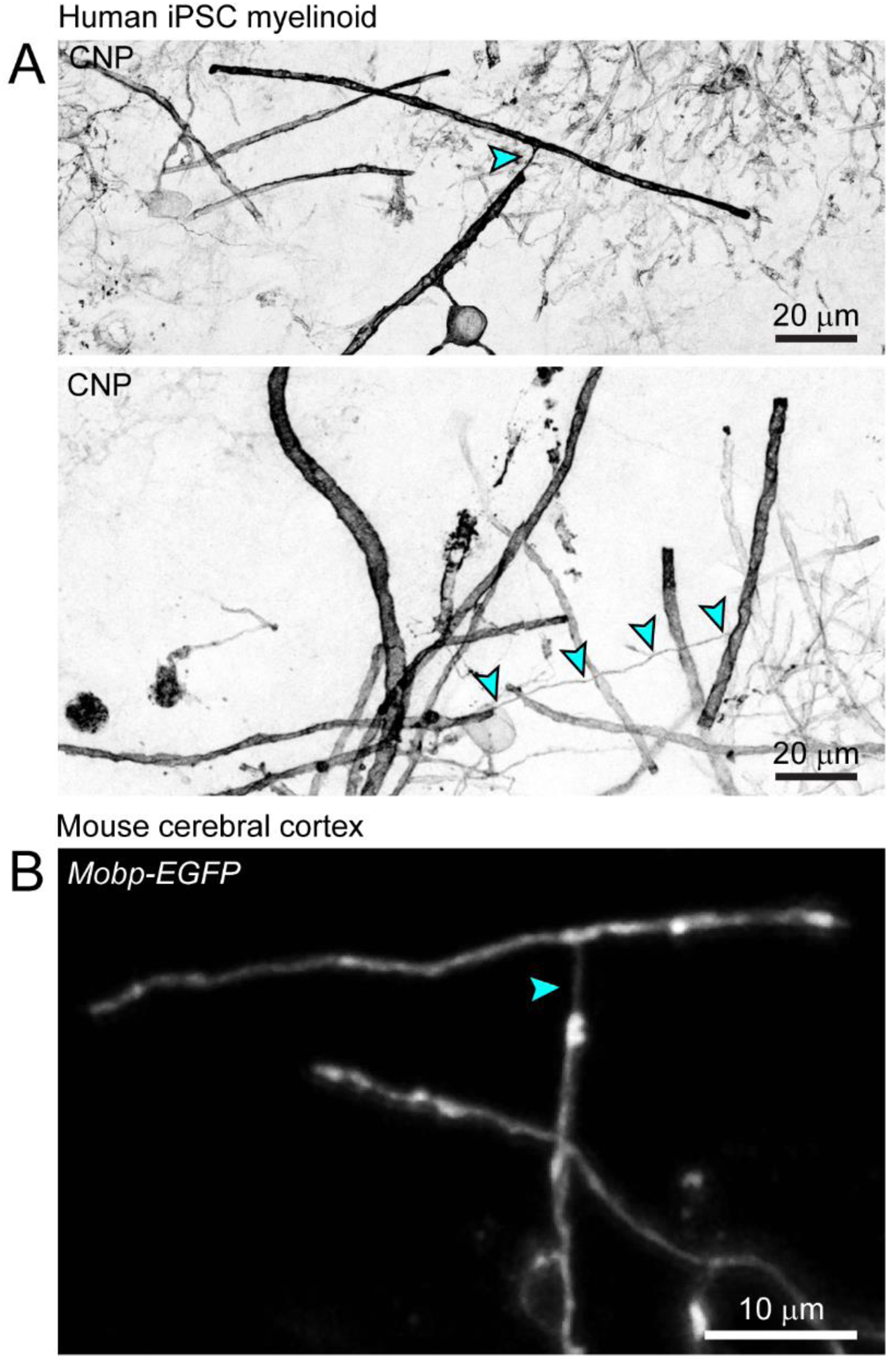
Unusual paranodal bridges. (A) Examples of atypical bridges connecting a paranode to an internode on different axons in human myelinoids.
(B) Example of an atypical paranode-internode bridge in the mouse cerebral cortex.

**Supplemental Figure 4:**
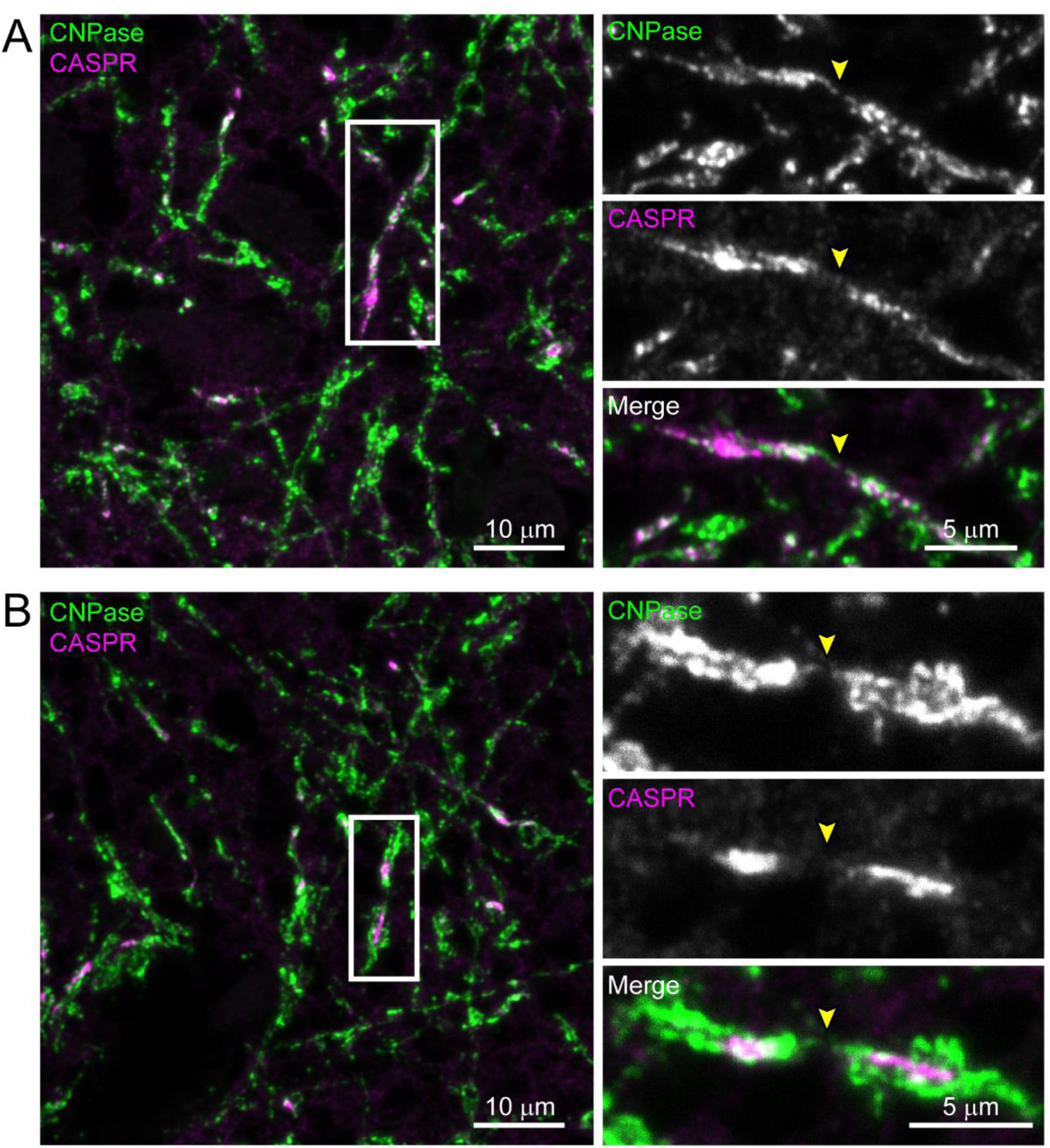
Paranodal bridges in human postmortem cerebral cortex. (A-B) Representative images of paranodal bridges in layer II/III of human post mortem motor cortex immunostained for CNPase and CASPR. Boxed region is magnified in separated channel images on the right. Arrowheads indicate CNPase-positive bridges spanning CASPR-positive paranodes.

**Supplemental Figure 5.**
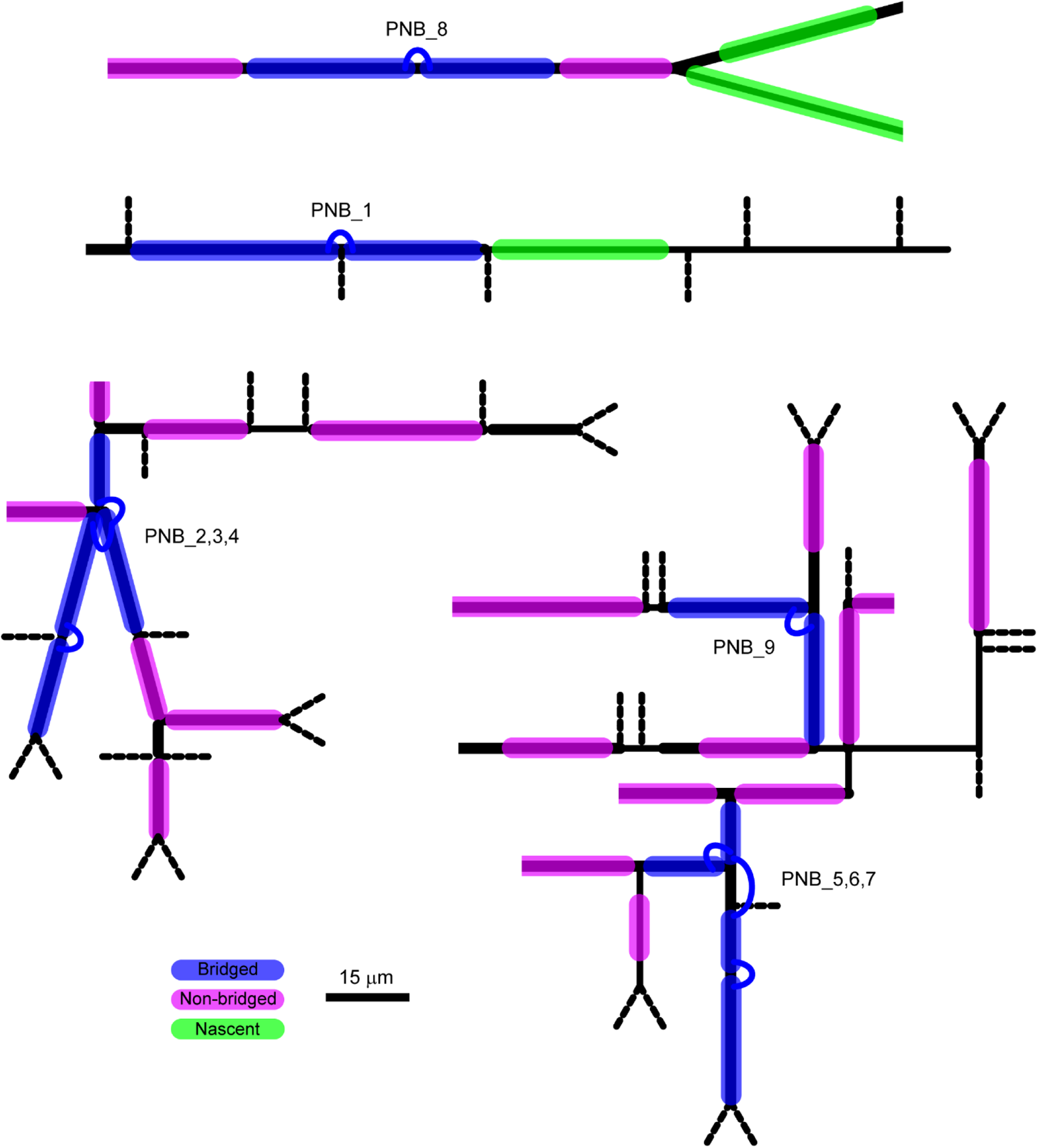
Skeletons of axonal morphologies myelinated by bridged sheaths. Skeletonized morphologies of four axons (black) with paranodal bridges in the EM dataset (Dorkenwald et al., 2019). Blue sheaths are those connected *via* paranodal bridges (connecting blue loops). Magenta sheaths are mature myelin sheaths without bridges with a direct cytoplasmic process. Green sheaths are non-compacted myelin sheaths presumably formed by recently differentiated oligodendrocytes. Axon segment lengths are scaled relative to the scale bar. Axon line thicknesses represent relative differences in diameter, but are not to scale. Dotted axon segments are very thin branches lined with synaptic terminals (full lengths not shown).

**Supplemental Figure 6.**
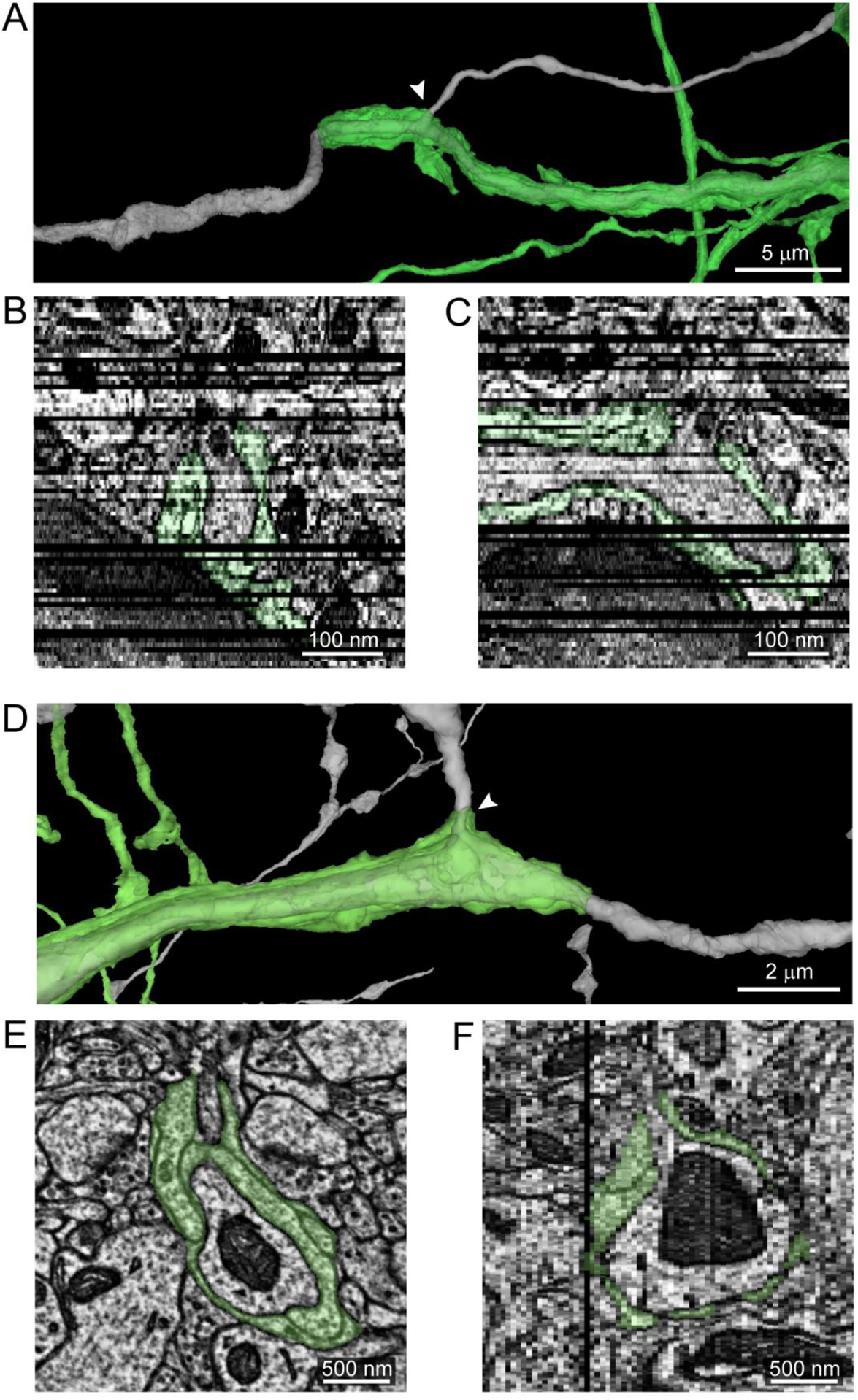
Nascent sheaths wrap axons across branch points prior to compaction. (A) Reconstructions of an axon (gray) and the processes of an immature oligodendrocyte that has generated sheaths, but none of them are compacted. One such nascent sheath wraps the axon past a branch point (arrowhead). Data from (Dorkenwald et al., 2019) and reconstructed with Neuroglancer (https://neuroglancer-demo.appspot.com/).
(B) XZ view of the position indicated by the arrowhead in *A*. Z slice depth 40 nm for *B* and *C*. Black lines are missing frames.
(C) YZ view of the position indicated by the arrowhead in *A*.
(D) Second example of a nascent sheath (green) crossing the branch point (arrowhead) of an axon (gray).
(E) XY view near the position indicated by the arrowhead in *D*.
(F) YZ view at the branch point of the axon in *D*.

**Supplemental Figure 7.**
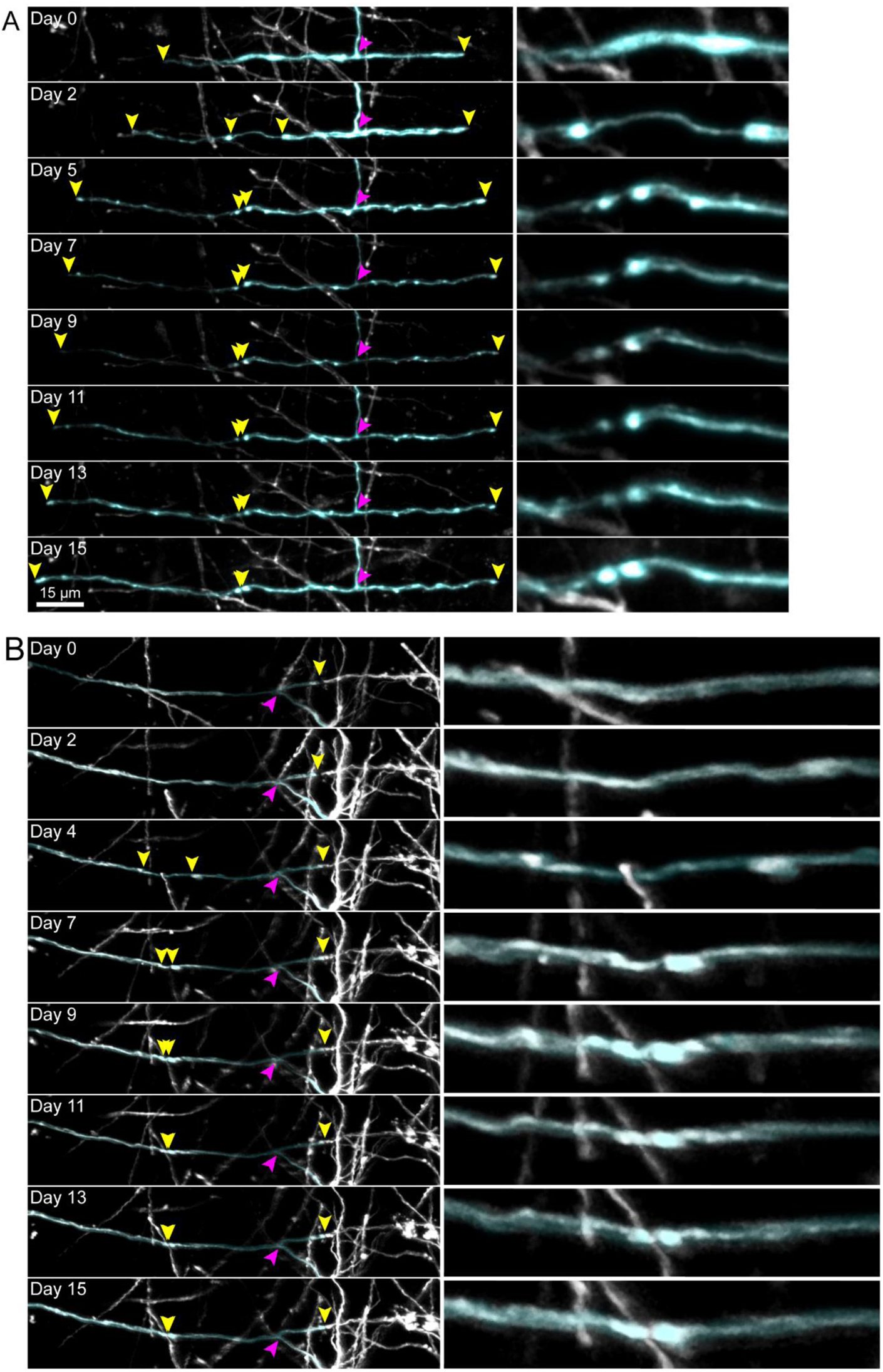
Individual nascent sheaths split, forming two sheaths connected by paranodal bridge across a node of Ranvier. (A) Time course of the formation of a mouse cortical paranodal bridge *in vivo*. At day 0, the oligodendrocyte has first appeared, and the highlighted sheath appears to begin as a single sheath. The magenta arrowhead indicates the intersection of the cell’s cytoplasmic process with this nascent sheath, and yellow arrowheads indicate the ends of this sheath. By day 2, the center of the sheath is thinned, and a thin strand of cytoplasm connects two bright puncta, which eventually form a doublet characteristic of paranodes flanking a node of Ranvier, as the paranodes move towards each other over the course of 15 days. The right column of images shows a magnified view of the forming paranodal bridge for the respective timepoint on the left.
(B) Time course of the formation of a second paranodal bridge *in vivo*.

**Supplemental Figure 8.**
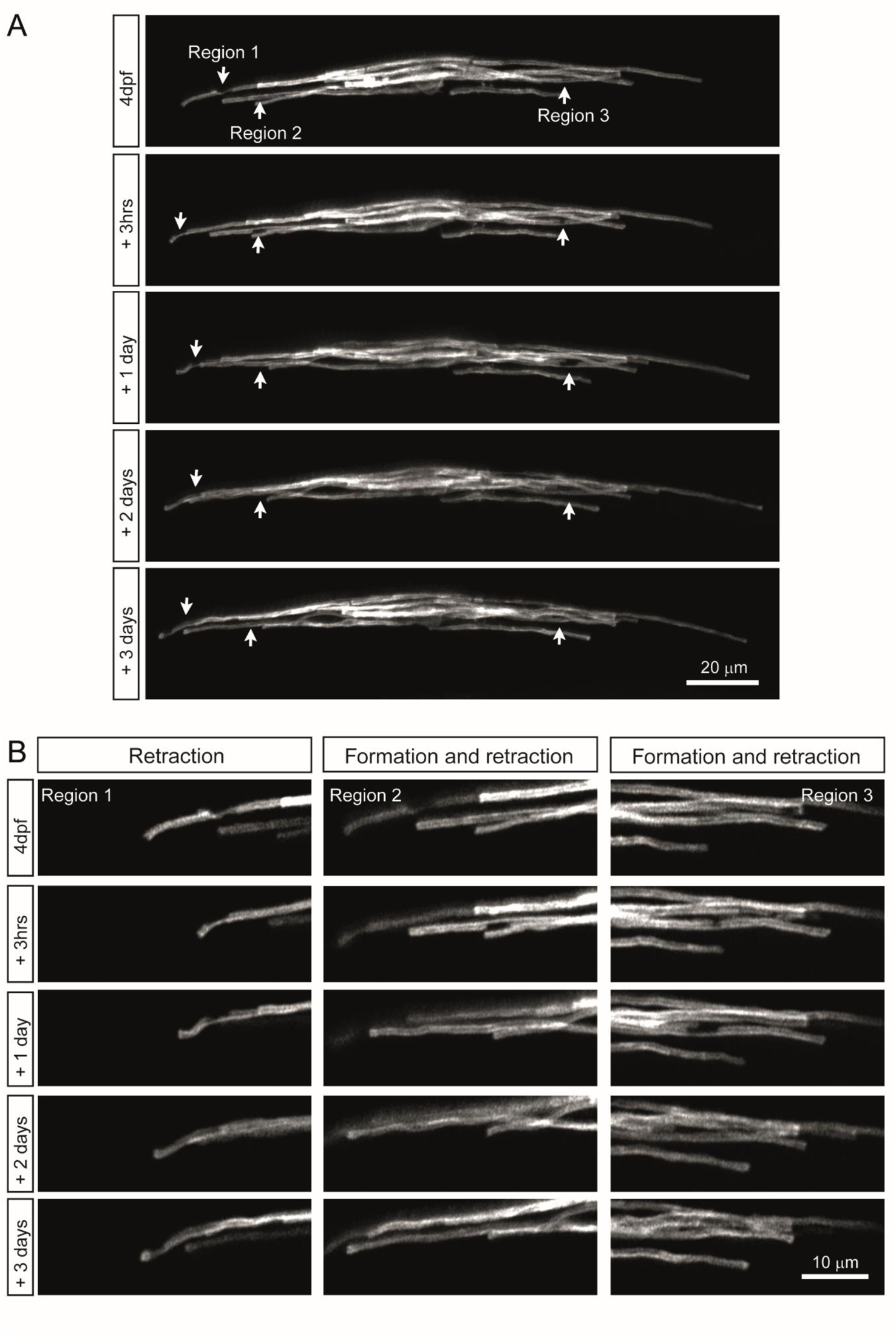
Dynamic paranodal bridges form by sheath splitting in the zebrafish spinal cord. (A) Time course of a single oligodendrocyte and its myelin sheaths (fluorescently labelled by mosaic expression of *mbp:EGFP-CAAX*) imaged between 4 and 7 dpf. Region 1, 2 and 3 show the locations where paranodal bridges form.
(B) Zoomed panels of regions 1, 2 and 3. Region 1 shows a myelin sheath which is present at 4 dpf but retracts by 7 dpf. Region 2 shows a myelin bridge forming between 4 dpf and + 3hrs which is no longer present by +1 day. Region 3 shows a myelin bridge forming between 4 dpf and + 3 hrs which is no longer present by +1 day.

## SUPPLEMENTAL MOVIES

**Supplemental Movie 1. Volumetric EM of paranodal bridge across an axon branch point.**

Continuous cytoplasm within outer tongues of two bridged sheaths highlighted magenta. Arrowheads track continuation of outer tongue cytoplasm across the paranodal bridge. Other paranodal loops are highlighted in cyan. Movie scrolls down through z stack before reversing back to top.

## REFERENCES

Aboul-Enein F, Rauschka H, Kornek B, Stadelmann C, Stefferl A, Brück W, Lucchinetti C, Schmidbauer M, Jellinger K, Lassmann H. 2003. Preferential Loss of Myelin-Associated Glycoprotein Reflects Hypoxia-Like White Matter Damage in Stroke and Inflammatory Brain Diseases. J Neuropathol Exp Neurol 62:25–33. doi:10.1093/jnen/62.1.25

Almeida RG, Czopka T, ffrench-Constant C, Lyons DA. 2011. Individual axons regulate the myelinating potential of single oligodendrocytes in vivo. Development 138:4443–4450. doi:10.1242/dev.071001

Almeida RG, Pan S, Cole KLH, Williamson JM, Early JJ, Czopka T, Klingseisen A, Chan JR, Lyons DA. 2018. Myelination of Neuronal Cell Bodies when Myelin Supply Exceeds Axonal Demand. Curr Biol 28:1296–1305.e5. doi:10.1016/j.cub.2018.02.068

Auer F, Vagionitis S, Czopka T. 2018. Evidence for Myelin Sheath Remodeling in the CNS Revealed by In Vivo Imaging. Curr Biol 28:549–559.e3. doi:10.1016/j.cub.2018.01.017

Bacmeister CM, Barr HJ, McClain CR, Thornton MA, Nettles D, Welle CG, Hughes EG. 2020. Motor learning promotes remyelination via new and surviving oligodendrocytes. Nat Neurosci 23:819–831. doi:10.1038/s41593-020-0637-3

Bechler ME, Byrne L, Ffrench-Constant C. 2015. CNS Myelin Sheath Lengths Are an Intrinsic Property of Oligodendrocytes. Curr Biol 25:2411–2416. doi:10.1016/j.cub.2015.07.056

Benveniste H, Liu X, Koundal S, Sanggaard S, Lee H, Wardlaw J. 2019. The Glymphatic System and Waste Clearance with Brain Aging: A Review. Gerontology 65:106–119. doi:10.1159/000490349

Boullerne AI. 2016. The history of myelin. Exp Neurol 283:431–445. doi:10.1016/j.expneurol.2016.06.005

Buchanan J, Elabbady L, Collman F, Jorstad NL, Bakken TE, Ott C, Glatzer J, Bleckert AA, Bodor AL, Brittan D, Bumbarger DJ, Mahalingam G, Seshamani S, Schneider-Mizell C, Takeno MM, Torres R, Yin W, Hodge RD, Castro M, Dorkenwald S, Ih D, Jordan CS, Kemnitz N, Lee K, Lu R, Macrina T, Mu S, Popovych S, Silversmith WM, Tartavull I, Turner NL, Wilson AM, Wong W, Wu J, Zlateski A, Zung J, Lippincott-Schwartz J, Lein ES, Seung HS, Bergles DE, Reid RC, Costa NM da. 2021. Oligodendrocyte precursor cells prune axons in the mouse neocortex. bioRxiv 2021.05.29.446047. doi:10.1101/2021.05.29.446047

Butt AM, Ransom BR. 1993. Morphology of astrocytes and oligodendrocytes during development in the intact rat optic nerve. J Comp Neurol 338:141–158. doi:10.1002/cne.903380110

Cai J, Qi Y, Hu X, Tan M, Liu Z, Zhang J, Li Q, Sander M, Qiu M. 2005. Generation of Oligodendrocyte Precursor Cells from Mouse Dorsal Spinal Cord Independent of Nkx6 Regulation and Shh Signaling. Neuron 45:41–53. doi:10.1016/j.neuron.2004.12.028

Call CL, Bergles DE. 2021. Cortical neurons exhibit diverse myelination patterns that scale between mouse brain regions and regenerate after demyelination. Nat Commun 12:4767. doi:10.1038/s41467-021-25035-2

Camandola S, Mattson MP. 2017. Brain metabolism in health, aging, and neurodegeneration. EMBO J 36:1474–1492. doi:10.15252/embj.201695810

Cardin JA, Carlén M, Meletis K, Knoblich U, Zhang F, Deisseroth K, Tsai L-H, Moore CI. 2009. Driving fast-spiking cells induces gamma rhythm and controls sensory responses. Nature 459:663–667. doi:10.1038/nature08002

Chong SYC, Rosenberg SS, Fancy SPJ, Zhao C, Shen Y-AA, Hahn AT, McGee AW, Xu X, Zheng B, Zhang LI, Rowitch DH, Franklin RJM, Lu QR, Chan JR. 2012. Neurite outgrowth inhibitor Nogo-A establishes spatial segregation and extent of oligodendrocyte myelination. Proc Natl Acad Sci 109:1299–1304. doi:10.1073/pnas.1113540109

Cobb SR, Buhl EH, Halasy K, Paulsen O, Somogyi P. 1995. Synchronization of neuronal activity in hippocampus by individual GABAergic interneurons. Nature 378:75–78. doi:10.1038/378075a0

Coleman M. 2005. Axon degeneration mechanisms: Commonality amid diversity. Nat Rev Neurosci. doi:10.1038/nrn1788

Czopka T, ffrench-Constant C, Lyons DA. 2013. Individual Oligodendrocytes Have Only a Few Hours in which to Generate New Myelin Sheaths In Vivo. Dev Cell 25:599–609. doi:10.1016/j.devcel.2013.05.013

del Rio Hortega P. 1922. ¿Son homologables la glia de escasas radiaciones y la celula de schwann? Boletín la Soc española Biol 10:25–28.

Dorkenwald S, Turner N, Macrina T, Lee K, Lu R, Wu J, Bodor A, Bleckert A, Brittain D, Kemnitz N, Silversmith W, Ih D, Zung J, Zlateski A, Tartavull I, Yu S-C, Popovych S, Wong W, Castro M, Jordan C, Wilson A, Froudarakis E, Buchanan J, Takeno M, Torres R, Mahalingam G, Collman F, Schneider-Mizell C, Bumbarger D, Li Y, Becker L, Suckow S, Reimer J, Tolias A, da Costa NM, Reid RC, Seung HS. 2019. Binary and analog variation of synapses between cortical pyramidal neurons. bioRxiv 2019.12.29.890319. doi:10.1101/2019.12.29.890319

Fogarty M, Richardson WD, Kessaris N. 2005. A subset of oligodendrocytes generated from radial glia in the dorsal spinal cord. Development 132:1951–1959. doi:10.1242/dev.01777

Geren B Ben. 1954. The formation from the schwann cell surface of myelin in the peripheral nerves of chick embryos. Exp Cell Res 7:558–562. doi:10.1016/S0014-4827(54)80098-X

Grossman Y, Parnas I, Spira ME. 1979a. Differential conduction block in branches of a bifurcating axon. J Physiol 295:283–305. doi:10.1113/JPHYSIOL.1979.SP012969

Grossman Y, Parnas I, Spira ME. 1979b. Mechanisms involved in differential conduction of potentials at high frequency in a branching axon. J Physiol 295:307–322. doi:10.1113/jphysiol.1979.sp012970

Haber M, Vautrin S, Fry EJ, Murai KK. 2009. Subtype-specific oligodendrocyte dynamics in organotypic culture. Glia 57:1000–1013. doi:10.1002/glia.20824

Hagemeyer N, Goebbels S, Papiol S, Kästner A, Hofer S, Begemann M, Gerwig UC, Boretius S, Wieser GL, Ronnenberg A, Gurvich A, Heckers SH, Frahm J, Nave KA, Ehrenreich H. 2012. A myelin gene causative of a catatonia-depression syndrome upon aging. EMBO Mol Med 4:528–539. doi:10.1002/emmm.201200230

Hamada MS, Kole MHP. 2015. Myelin Loss and Axonal Ion Channel Adaptations Associated with Gray Matter Neuronal Hyperexcitability. J Neurosci 35:7272–7286. doi:10.1523/JNEUROSCI.4747-14.2015

Hardy RJ, Friedrich VL. 1996. Progressive remodeling of the oligodendrocyte process arbor during myelinogenesis. Dev Neurosci 18:243–254. doi:10.1159/000111414

Harris JJ, Attwell D. 2012. The Energetics of CNS White Matter. J Neurosci 32:356–371. doi:10.1523/JNEUROSCI.3430-11.2012

Hill RA, Li AM, Grutzendler J. 2018. Lifelong cortical myelin plasticity and age-related degeneration in the live mammalian brain. Nat Neurosci 21:683–695. doi:10.1038/s41593-018-0120-6

Hughes AN, Appel B. 2020. Microglia phagocytose myelin sheaths to modify developmental myelination. Nat Neurosci. doi:10.1038/s41593-020-0654-2

Hughes EG, Kang SH, Fukaya M, Bergles DE. 2013. Oligodendrocyte progenitors balance growth with self-repulsion to achieve homeostasis in the adult brain. Nat Neurosci 16:668– 676. doi:10.1038/nn.3390

Hughes EG, Orthmann-Murphy JL, Langseth AJ, Bergles DE. 2018. Myelin remodeling through experience-dependent oligodendrogenesis in the adult somatosensory cortex. Nat Neurosci 21:696–706. doi:10.1038/s41593-018-0121-5

Ioannidou K, Anderson KI, Strachan D, Edgar JM, Barnett SC. 2012. Time-lapse imaging of the dynamics of CNS glial-axonal interactions in vitro and ex vivo. PLoS One 7. doi:10.1371/journal.pone.0030775

James OG, Selvaraj BT, Magnani D, Burr K, Connick P, Barton SK, Vasistha NA, Hampton DW, Story D, Smigiel R, Ploski R, Brophy PJ, Ffrench-Constant C, Lyons DA, Chandran S. 2021. iPSC-derived myelinoids to study myelin biology of humans. Dev Cell 56:1346–1358.e6. doi:10.1016/j.devcel.2021.04.006

Kessaris N, Fogarty M, Iannarelli P, Grist M, Wegner M, Richardson WD. 2006. Competing waves of oligodendrocytes in the forebrain and postnatal elimination of an embryonic lineage. Nat Neurosci 9:173–179. doi:10.1038/nn1620

Kirby BB, Takada N, Latimer AJ, Shin J, Carney TJ, Kelsh RN, Appel B. 2006. In vivo time-lapse imaging shows dynamic oligodendrocyte progenitor behavior during zebrafish development. Nat Neurosci 9:1506–1511. doi:10.1038/nn1803

Klausberger T, Márton LF, Baude A, Roberts JDB, Magill PJ, Somogyi P. 2004. Spike timing of dendrite-targeting bistratified cells during hippocampal network oscillations in vivo. Nat Neurosci 7:41–47. doi:10.1038/nn1159

Koudelka S, Voas MG, Almeida RG, Baraban M, Soetaert J, Meyer MP, Talbot WS, Lyons DA. 2016. Individual Neuronal Subtypes Exhibit Diversity in CNS Myelination Mediated by Synaptic Vesicle Release. Curr Biol 26:1447–1455. doi:10.1016/j.cub.2016.03.070

Lang EJ, Rosenbluth J. 2003. Role of myelination in the development of a uniform olivocerebellar conduction time. J Neurophysiol 89:2259–70. doi:10.1152/jn.00922.2002

Lappe-Siefke C, Goebbels S, Gravel M, Nicksch E, Lee J, Braun PE, Griffiths IR, Nave K-A. 2003. Disruption of Cnp1 uncouples oligodendroglial functions in axonal support and myelination. doi:10.1038/ng1095

Larson VA, Mironova Y, Vanderpool KG, Waisman A, Rash JE, Agarwal A, Bergles DE. 2018. Oligodendrocytes control potassium accumulation in white matter and seizure susceptibility. Elife 7:1–33. doi:10.7554/eLife.34829

Lee Y, Morrison BM, Li Y, Lengacher S, Farah MH, Hoffman PN, Liu Y, Tsingalia A, Jin L, Zhang PW, Pellerin L, Magistretti PJ, Rothstein JD. 2012. Oligodendroglia metabolically support axons and contribute to neurodegeneration. Nature 487:443–448. doi:10.1038/nature11314

Manor Y, Koch C, Segev I. 1991. Effect of geometrical irregularities on propagation delay in axonal trees. Biophys J 60:1424–1437. doi:10.1016/S0006-3495(91)82179-8

Marisca R, Hoche T, Agirre E, Hoodless LJ, Barkey W, Auer F, Castelo-Branco G, Czopka T. 2020. Functionally distinct subgroups of oligodendrocyte precursor cells integrate neural activity and execute myelin formation. Nat Neurosci 23:363–374. doi:10.1038/s41593-019-0581-2

Mattson MP, Arumugam T V. 2018. Hallmarks of Brain Aging: Adaptive and Pathological Modification by Metabolic States. Cell Metab 27:1176–1199. doi:10.1016/J.CMET.2018.05.011

Mayoral SR, Etxeberria A, Shen Y-AA, Chan JR. 2018. Initiation of CNS Myelination in the Optic Nerve Is Dependent on Axon Caliber. Cell Rep 25:544–550.e3. doi:10.1016/j.celrep.2018.09.052

Micheva KD, Kiraly M, Perez MM, Madison D V. 2021. Conduction Velocity Along the Local Axons of Parvalbumin Interneurons Correlates With the Degree of Axonal Myelination. Cereb Cortex 31:3374–3392. doi:10.1093/CERCOR/BHAB018

Micheva KD, Wolman D, Mensh BD, Pax E, Buchanan J, Smith SJ, Bock DD. 2016. A large fraction of neocortical myelin ensheathes axons of local inhibitory neurons. Elife 5:1–29. doi:10.7554/eLife.15784

Morrison BM, Tsingalia A, Vidensky S, Lee Y, Jin L, Farah MH, Lengacher S, Magistretti PJ, Pellerin L, Rothstein JD. 2015. Deficiency in monocarboxylate transporter 1 (MCT1) in mice delays regeneration of peripheral nerves following sciatic nerve crush. Exp Neurol 263:325–338. doi:10.1016/j.expneurol.2014.10.018

Murtie JC, Macklin WB, Corfas G. 2007. Morphometric analysis of oligodendrocytes in the adult mouse frontal cortex. J Neurosci Res 85:2080–2086. doi:10.1002/jnr.21339

Orthmann-Murphy J, Call CL, Molina-Castro GC, Hsieh YC, Rasband MN, Calabresi PA, Bergles DE. 2020. Remyelination alters the pattern of myelin in the cerebral cortex. Elife 9:1–61. doi:10.7554/eLife.56621

Pajevic S, Basser PJ, Fields RD. 2014. Role of myelin plasticity in oscillations and synchrony of neuronal activity. Neuroscience 276:135–147. doi:10.1016/j.neuroscience.2013.11.007

Parnas I, Segev I. 1979. A mathematical model for conduction of action potentials along bifurcating axons. J Physiol 295:323–343. doi:10.1113/jphysiol.1979.sp012971

Penfield W. 1924. Oligodendroglia and its relation to classical neuroglia. Brain 47:430–452. doi:10.1093/brain/47.4.430

Philips T, Mironova YA, Jouroukhin Y, Chew J, Vidensky S, Farah MH, Pletnikov M V., Bergles DE, Morrison BM, Rothstein JD. 2021. MCT1 Deletion in Oligodendrocyte Lineage Cells Causes Late-Onset Hypomyelination and Axonal Degeneration. Cell Rep 34. doi:10.1016/j.celrep.2020.108610

Portera-Cailliau C, Weimer RM, De Paola V, Caroni P, Svoboda K. 2005. Diverse modes of axon elaboration in the developing neocortex. PLoS Biol 3. doi:10.1371/journal.pbio.0030272

Ratnadurai-Giridharann S, Khargonekar PP, Talathi SS. 2015. Emergent gamma synchrony in all-to-all interneuronal networks. Front Comput Neurosci 9. doi:10.3389/fncom.2015.00127

Rinholm JE, Hamilton NB, Kessaris N, Richardson WD, Bergersen LH, Attwell D. 2011. Regulation of Oligodendrocyte Development and Myelination by Glucose and Lactate. J Neurosci 31:538–548. doi:10.1523/JNEUROSCI.3516-10.2011

Saab AS, Nave KA. 2017. Myelin dynamics: protecting and shaping neuronal functions. Curr Opin Neurobiol 47:104–112. doi:10.1016/j.conb.2017.09.013

Saab AS, Tzvetanova ID, Nave KA. 2013. The role of myelin and oligodendrocytes in axonal energy metabolism. Curr Opin Neurobiol 23:1065–1072. doi:10.1016/j.conb.2013.09.008

Schirmer L, Möbius W, Zhao C, Cruz-Herranz A, Ben Haim L, Cordano C, Shiow LR, Kelley KW, Sadowski B, Timmons G, Pröbstel AK, Wright JN, Sin JH, Devereux M, Morrison DE, Chang SM, Sabeur K, Green AJ, Nave KA, Franklin RJM, Rowitch DH. 2018. Oligodendrocyte-encoded kir4.1 function is required for axonal integrity. Elife 7:1–21. doi:10.7554/eLife.36428

Schmitt FO, Bear RS, Clark GL. 1935. X-ray Diffraction Studies on Nerve. Radiology 25:131– 151. doi:10.1148/25.2.131

Segovia KN, McClure M, Moravec M, Luo NL, Wan Y, Gong X, Riddle A, Craig A, Struve J, Sherman LS, Back SA. 2008. Arrested oligodendrocyte lineage maturation in chronic perinatal white matter injury. Ann Neurol 63:520–530. doi:10.1002/ana.21359

Seidl AH, Rubel EW. 2016. Systematic and differential myelination of axon collaterals in the mammalian auditory brainstem. Glia 64:487–494. doi:10.1002/glia.22941

Seidl AH, Rubel EW, Barria A. 2014. Differential Conduction Velocity Regulation in Ipsilateral and Contralateral Collaterals Innervating Brainstem Coincidence Detector Neurons. J Neurosci 34:4914–4919. doi:10.1523/JNEUROSCI.5460-13.2014

Snaidero N, Möbius W, Czopka T, Hekking LHPHP, Mathisen C, Verkleij D, Goebbels S, Edgar J, Merkler D, Lyons DAA, Nave K-AA, Simons M. 2014. Myelin membrane wrapping of CNS axons by PI(3,4,5)P3-dependent polarized growth at the inner tongue. Cell 156:277–290. doi:10.1016/j.cell.2013.11.044

Snaidero N, Velte C, Myllykoski M, Raasakka A, Ignatev A, Werner HB, Erwig MS, Möbius W, Kursula P, Nave KA, Simons M. 2017. Antagonistic Functions of MBP and CNP Establish Cytosolic Channels in CNS Myelin. Cell Rep 18:314–323. doi:10.1016/j.celrep.2016.12.053

Stedehouder J, Brizee D, Shpak G, Kushner SA. 2018. Activity-Dependent Myelination of Parvalbumin Interneurons Mediated by Axonal Morphological Plasticity. J Neurosci 38:3631–3642. doi:10.1523/JNEUROSCI.0074-18.2018

Stedehouder J, Brizee D, Slotman JA, Pascual-García M, Leyrer ML, Bouwen BLJ, Dirven CMF, Gao Z, Berson DM, Houtsmuller AB, Kushner SA. 2019. Local axonal morphology guides the topography of interneuron myelination in mouse and human neocortex. Elife 8:1–28. doi:10.7554/eLife.48615

Stedehouder J, Couey JJ, Brizee D, Hosseini B, Slotman JA, Dirven CMFF, Shpak G, Houtsmuller AB, Kushner SA. 2017. Fast-spiking Parvalbumin Interneurons are Frequently Myelinated in the Cerebral Cortex of Mice and Humans. Cereb Cortex 27:5001–5013. doi:10.1093/cercor/bhx203

Stedehouder J, Kushner SA. 2017. Myelination of parvalbumin interneurons: A parsimonious locus of pathophysiological convergence in schizophrenia. Mol Psychiatry 22:4–12. doi:10.1038/mp.2016.147

Toth E, Rassul SM, Berry M, Fulton D. 2021. A morphological analysis of activity-dependent myelination and myelin injury in transitional oligodendrocytes. Sci Rep 11. doi:10.1038/s41598-021-88887-0

Tripathi RB, Jackiewicz M, McKenzie IA, Kougioumtzidou E, Grist M, Richardson WD. 2017. Remarkable Stability of Myelinating Oligodendrocytes in Mice. Cell Rep 21:316–323. doi:10.1016/j.celrep.2017.09.050

Uzor N-E, McCullough LD, Tsvetkov AS. 2020. Peroxisomal Dysfunction in Neurological Diseases and Brain Aging. Front Cell Neurosci 14:44. doi:10.3389/fncel.2020.00044

Vallstedt A, Klos JM, Ericson J. 2005. Multiple dorsoventral origins of oligodendrocyte generation in the spinal cord and hindbrain. Neuron 45:55–67. doi:10.1016/j.neuron.2004.12.026

van Leeuwenhoek A. 1719. Epistola XXXIIEpistolae Physiologicae Super Compluribus Naturae Arcanis. Delft, Adrianum Beman. pp. 309–317.

Wang F, Ren S-Y, Chen J-F, Liu K, Li R-X, Li Z-F, Hu B, Niu J-Q, Xiao L, Chan JR, Mei F. 2020. Myelin degeneration and diminished myelin renewal contribute to age-related deficits in memory. Nat Neurosci 2020 234 23:481–486. doi:10.1038/s41593-020-0588-8

Wang XJ, Buzsáki G. 1996. Gamma oscillation by synaptic inhibition in a hippocampal interneuronal network model. J Neurosci 16:6402–6413. doi:10.1523/jneurosci.16-20-06402.1996

Yamahachi H, Marik SA, McManus JNJ, Denk W, Gilbert CD. 2009. Rapid Axonal Sprouting and Pruning Accompany Functional Reorganization in Primary Visual Cortex. Neuron 64:719–729. doi:10.1016/J.NEURON.2009.11.026

Yeung MSYSY, Zdunek S, Bergmann O, Bernard S, Salehpour M, Alkass K, Perl S, Tisdale J, Possnert G, Brundin L, Druid H, Frisén J. 2014. Dynamics of Oligodendrocyte Generation and Myelination in the Human Brain. Cell 159:766–774. doi:10.1016/j.cell.2014.10.011

Young KM, Psachoulia K, Tripathi RB, Dunn SJ, Cossell L, Attwell D, Tohyama K, Richardson WD. 2013. Oligodendrocyte dynamics in the healthy adult CNS: Evidence for myelin remodeling. Neuron 77:873–885. doi:10.1016/j.neuron.2013.01.006

Ziabreva I, Campbell G, Rist J, Zambonin J, Rorbach J, Wydro MM, Lassmann H, Franklin RJM, Mahad D. 2010. Injury and differentiation following inhibition of mitochondrial respiratory chain complex IV in rat oligodendrocytes. Glia 58:1827–1837. doi:10.1002/glia.21052

Zonouzi M, Berger D, Jokhi V, Kedaigle A, Lichtman J, Arlotta P. 2019. Individual Oligodendrocytes Show Bias for Inhibitory Axons in the Neocortex. Cell Rep 27:2799–2808.e3. doi:10.1016/j.celrep.2019.05.018

